# A neurocomputational account of the link between social perception and social action

**DOI:** 10.1101/2023.10.02.560256

**Authors:** Lisa M. Bas, Ian D. Roberts, Cendri A. Hutcherson, Anita Tusche

**Author notes:** Corresponding author: Dr. Lisa M. Bas, Queen’s University, 99 University Ave, Kingston, ON K7L 3N6. Authors contributed equally (shared first authorship) (Lisa M. Bas, Ian D. Roberts). Authors contributed equally (shared last authorship) (Cendri A. Hutcherson, Anita Tusche).

## Abstract

People selectively help others based on perceptions of their merit or need. Here, we develop a neurocomputational account of how these social perceptions translate into social choice. Using a novel fMRI social perception task, we show that both merit and need perceptions recruited the brain’s social inference network. A behavioral computational model identified two non-exclusive mechanisms underlying variance in social perceptions: a consistent tendency to perceive others as meritorious/needy (bias) and a propensity to sample and integrate normative evidence distinguishing high from low merit/need in other people (sensitivity). Variance in people’s merit (but not need) bias and sensitivity independently predicted distinct aspects of altruism in a social choice task completed months later. An individual’s merit *bias* predicted *context-independent* variance in people’s overall other-regard during altruistic choice, biasing people towards prosocial actions. An individual’s merit *sensitivity* predicted *context-sensitive* discrimination in generosity towards high and low merit recipients by influencing other-regard and self-regard during altruistic decision-making. This context-sensitive perception-action link was associated with activation in the right temporoparietal junction. Together, these findings point towards stable, biologically based individual differences in perceptual processes related to abstract social concepts like merit, and suggest that these differences may have important behavioral implications for an individual’s tendency toward favoritism or discrimination in social settings.

## Introduction

Psychologists and economists have long sought to explain when and why people help. While people are generally altruistic, they also show selectivity, being more likely to assist those in need ^1–5^ and to withhold aid from those perceived as undeserving ^6–10^. Need and deservingness (merit) are two distinct principles of morality. The need principle involves distributing resources to those who require them, irrespective of whether they have earned them, while the “merit principle” focuses on allocating resources based on individuals’ deservingness, regardless of their actual need ^11^. How do people assess whether others deserve or need help, and how does this influence their choices to initiate that help?

Based on an analogy to perception in basic sensory domains like vision ^12, 13^, we hypothesized that variance in social perceptions might be driven by two discrete mechanisms: an individual’s *sensitivity* to social situational cues signaling other’s merit or need, and an individual’s *bias* to perceive merit or need independently of specific cues. In other words, one can think of the likelihood *P* that person *J* perceives a target individual as deserving (meritorious) or in need of aid as being determined by the sum of all cues *C_1_, C_2_, … C_N_* associated with that perceptual judgment, multiplied by a person’s idiosyncratic sensitivity *S* to each cue, and added to their baseline tendency (*bias*) to perceive others as deserving or in need:

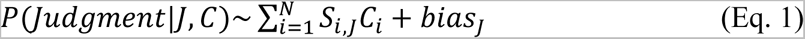

Need-signaling cues could include facial, vocal, or postural cues of pain or distress ^14–17^, or cues implying imminent harm (e.g., a person in front of a runaway car ^18, 19^). Merit-relevant cues could include membership in ‘good’ or ‘bad’ groups (e.g., children ^20^, one’s in-group ^21^, Nazi Party membership ^22^) or information about the social normativity or benevolence of that person’s actions^23^. An individual with higher average *sensitivity* to the cues associated with that judgment would discriminate more between those who are high or low in merit or need (as signaled by available cues). An individual with a larger *bias* term would simply be more likely to judge *all* individuals as meritorious/needy irrespective of present cues. Whether processing and integrating these social cues, along with the resulting judgments, involves distinct neural modules ^24^ or a general-purpose mechanism for social cognition ^25^, such as the mentalizing network ^26–29^, remains unknown.

A similar question arises over *how* perceptions of need and merit influence meaningful *behavior*. While extensive research shows that they do ^3–6, 8–10, 30–32^, we know little about the precise computational mechanism underlying this perception-action link. We speculated that it might operate by changing the value that people ascribe to different social considerations during choice. More specifically, we build on a growing body of work suggesting that prosocial actions can be characterized as a value-based decision process ^33–36^. In this framework, the value of acting prosocial is represented as a weighted sum of several attributes, like the cost of helping for the decision-maker, the benefits for others, or the fairness of the outcome. These three attributes have been repeatedly shown to guide social behaviors ^37–43^, with higher weights on prosocial attributes like benefits for others or fairness yielding more prosocial choices ^35, 44–49^. Within this computational framework, perceptions of others’ merit or need could affect social behaviors through a simple mechanism: by altering the weights given to outcomes for self, outcomes for others, or fairness during the decision process. For example, perceiving someone as highly deserving or needy could increase helping by increasing the weight given to the recipient’s benefits, decreasing the weight given to one’s own benefits, or both.

This computational framework leads to a set of testable predictions about how individual differences in social perceptual sensitivity and bias (Eq. 1) might influence altruistic behavior: An individual’s general tendency to perceive merit or need (i.e., their *bias* parameter) should be correlated with a general tendency to ascribe higher weight to prosocial considerations (i.e., others’ benefits or fairness). In contrast, an individual’s sensitivity to cues signaling merit or need (i.e., their average *sensitivity* parameter *S*) should make them more discriminatory, increasing weights on prosocial considerations for those judged as deserving (or needy) but decreasing them for those judged as lacking these qualities. These weights should then manifest in the frequency with which an individual acts generously, either overall or as a function of the recipient’s merit and/or need.

To test these hypotheses, and to uncover their neural basis, we first asked participants to complete a novel social perception task while we collected their brain responses using fMRI (functional magnetic resonance imaging). The behavior observed in this task allowed us to computationally estimate two distinct – but not mutually exclusive – processes underlying individual differences in merit/need perception: an overall propensity to perceive others as deserving/in need (i.e., an individual’s *bias* parameter) and a tendency to sample and integrate cues signaling merit/need (i.e., an individual’s *sensitivity* parameter). Neurally, we investigated whether merit and need are perceived through distinct or common neural circuits and reflect individual differences in people’s perceptual bias and/or sensitivity. Examining need and merit concurrently in this task will also help clarify the computational and neural underpinnings of these related, but distinct concepts, distinguishing between them more effectively. Second, we used a separate altruistic choice task (completed on average ∼303 days later) to computationally estimate the weights people place on themselves, others, and fairness when deciding whether to provide aid, and we examined how those weights varied as a function of recipient merit and need. Third, we tested whether individual differences in merit/need bias or sensitivity and their neural underpinnings in the social perception task predict individual differences in prosocial action in the altruism task. Our results suggest that merit (but perhaps not need) perceptions result from stable individual differences in both bias and sensitivity and manifest in an individual’s prosocial behavior months later.

## Materials and Methods

### Participants

This research took place in the context of a large project that recruited participants from the broader Los Angeles metropolitan area to come to the lab on year> tag found, Please Check. ln:357 cl:32 [Edit].days to complete different behavioral and neuroimaging tasks related to social cognition ^50^. We recruited 50 participants from this larger project pool to complete a newly developed fMRI social perception task. All participants were right-handed, had normal or corrected-to-normal vision, spoke English fluently, and had IQ scores in the normal range (20 females; mean age = 33 years, range = 19–49; full-scale IQ = 105.18 ± 8.04 (mean ± std), range = 87–127 ^51, 52^). Of these 50 participants, we excluded one who fell asleep and five for excessive movement (framewise displacement > 0.3 mm on over 30% of frames and visual spikes), yielding a total sample of n = 44 for this task. We also recruited 42 participants from the larger participant pool to complete the altruism task (13 females; mean age = 34 years, range = 19–49; full-scale IQ = 104.79 ± 7.97, range = 87–127). Of these 42 participants, we excluded one individual who fell asleep, five for excessive movement during fMRI data collection, three for invariant behavioral responses (identical left/right button press in > 90% of trials, indicating inattention to monetary offers in the altruism task), and five for manipulation check failures (e.g., failing memory checks regarding partner behavior in the altruism task, see below). This procedure yielded a sample of n = 28 for the altruism task, with an overlap of 25 individuals who successfully completed both sessions (social perception and altruism task). Thus, we report results regarding the social perception task for 44 participants (27 males; mean age = 34 years, range = 19–49; full-scale IQ = 105.84 ± 7.85, range = 90–127), the altruism task for 28 participants (19 males; mean age = 35 years, range = 25–49; full-scale IQ = 106.36 ± 7.94, range = 94–127), and cross-session results (comparing data across both tasks) for the sample of 25 individuals with valid data in both (17 males; mean age = 35 years, range = 25–49; full-scale IQ = 106.60 ± 8.26, range = 94–127). Participants in our overlap sample (valid data in both tasks) had an average separation between both tasks of 303 days (range: 27-663). This delay minimized the risk of sequential dependencies between tasks and increased confidence in the temporal stability of relationships between social perception and social action. Participants received $20 per hour for each experimental session and an additional amount based on one randomly selected trial in the altruism task to incentivize choices and ensure that participants’ responses reflected their actual preferences. All participants provided written informed consent according to a protocol approved by the Institutional Review Board of the California Institute of Technology (#12-0343).

### Social Perception Task

To assess individual differences in social perceptions, we used a modified version of the established fMRI why/how task ^53, 54^. Participants viewed images of people in complex real-world scenes and made rapid yes/no judgments (button presses) while their brain responses were measured using fMRI. In separate blocks (Figure 1A), participants made judgments regarding others’ perceived merit (“Does this person deserve help?”), need (“Does this person need help?”), or factual judgments that did not require social inferences (“Does this person use both hands?”), which served as a control condition. Before each task block, a visual prompt informed participants of the upcoming condition of the task block. Moreover, a keyword was briefly presented between images of a block as a reminder (i.e., “need”, “deserve”, or “both hands”; Figure 1A). Each block consisted of 32 images, and participants completed two blocks per condition. Thus, participants viewed a total of 64 images, each presented once per condition (need, merit, and control). Some images displayed year> tag found, Please Check. ln:357 cl:32 [Edit].people; a green arrow superimposed on the photograph indicated the target of the social perception. Images were displayed for 2 seconds with a 0.5-second inter-stimulus interval. The presentation order of images was fixed across participants to maximize efficiency. Crucially, this task did not require people to make altruistic choices or engage in meaningful social interaction. Instead, it focused solely on capturing individuals’ patterns of perceiving others’ perceived merit, need, and factual control inferences, which we modeled using an evidence accumulation framework (computational model of social perception, see below). The task was implemented in MATLAB (MathWorks) using the Psychophysics Toolbox extensions ^55–57^. The stimuli and presentation code are available at https://osf.io/4u5vs/.

**Figure 1.**
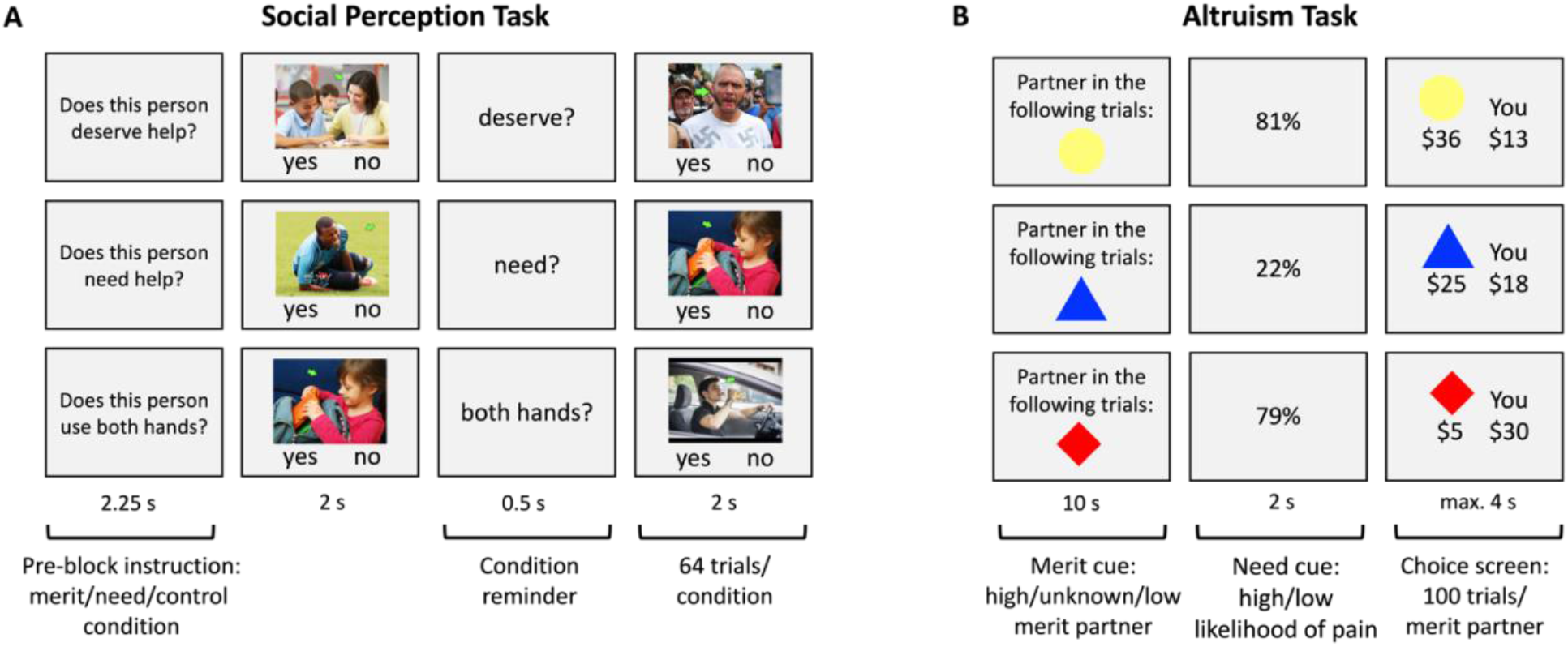
Experimental tasks performed on separate days. (**A)** Social Perception Task. Participants made rapid yes/no judgments regarding others’ perceived deservingness (merit blocks, top row), need (need blocks, middle row), or factual inferences (control blocks, bottom row) while their brain responses were measured using fMRI. (**B**) Altruism Task. On each trial, participants accepted or rejected a monetary offer that affected the payoffs for themselves (“You”) and one of three partners (choice screen; displayed offer vs. constant default of $20 for both). The three partners (identified via colored geometric shapes) differed in their associated merit (merit cue: high/low/unknown) based on partner behavior in a separate exchange game played before the altruism task. Partner’s need (need cue: high/low) was manipulated on a trial-by-trial basis, indicated by the likelihood of a painful cold pressor task (CPT) for the partner after the altruism task (high need: 80±4%; low need: 20±4%). Partners could buy out of the painful CPT using funds from one randomly selected trial at the end of the altruism task. Participants were informed that they could help their partners to avoid the painful CPT by making generous choices.

### Altruism Task

Participants completed an altruism task (modified dictator game) on a different day than the social perception task (average delay of 303 days, min. = 27, max. = 663). All but one participant completed the social perception task first. On each trial, we presented participants with a proposed monetary allocation between themselves and one of three partners (e.g., $15 for themselves and $25 for their current partner; choice screen, see Figure 1B). Participants decided whether to accept or reject the proposed allocation in favor of a constant default allocation of $20 for both ^45, 58^. Participants indicated their choices by pressing one of four buttons (“strong no”, “no”, “yes”, or “strong yes”). The direction of the response scale (“strong yes” to “strong no”) and the presentation side of self- and other-related payoffs (left vs. right side of the screen) was counterbalanced across participants. Proposed monetary outcomes for the participant (“You”) and the partner (represented by one of three colored geometric shapes) ranged from $5 to $35 (see Supplemental Figure S1). To minimize habituation and repetition effects, we randomly jittered proposal amounts by $0-$4. We informed participants that one trial would be randomly selected and implemented at the end of the experiment. In total, there were 300 trials divided across five runs (i.e., 60 trials per run, 100 trials per partner). Stimulus presentation and response collection in the altruism task were implemented using PsychoPy ^59, 60^.

*Partner’s Merit (High/Low/Unknown).* Participants played the altruism task with three partners that differed in their perceived merit (implemented in separate blocks; 20 consecutive trials per partner within a functional run; counterbalanced order of the three partners across participants). The merit of the three partners in the task was manipulated before the altruism task and was based on information about partners’ behavior in a sequential prisoners’ dilemma task that partners played with anonymous third persons, modified from ^61, 62^ (see Supplemental Note S1). Merit levels were manipulated such that one player was perceived as highly deserving (high merit partner), undeserving (low merit partner), or having unknown merit (no information provided before the altruism task, control condition). Partner identity was indicated in the altruism task using one of three colored geometric shapes (random combination of either a circle, diamond, or triangle, colored either red, yellow, or blue), shown at the beginning of each block of 20 trials with the same partner (merit cue, Figure 1B) and on each trial-wise offer screen indicating payoffs for the specific partner (choice screen, Figure 1B).

*Partner’s Need (High/Low).* The altruism task also experimentally manipulated the need level of the three partners on a trial-by-trial level. Specifically, we manipulated the probability that a partner would have to complete a painful cold pressor task (CPT ^63^) at the end of the experiment (outside of the scanner; hand submerged in ice water for ∼2 minutes). We informed participants that each of the three partners would be given the option of using money received in the altruism task to buy out of the post-task CPT (based on a randomly selected trial at the end of the task that would be implemented according to the participants’ choice on that trial). With each dollar spent, the partner could subtract 10% from their probability of having to perform the painful CPT (e.g., spending $3 would reduce an 80% chance to 50%). Thus, participants knew they could help the other player avoid the painful CPT by making generous choices.

To signal need on each trial, participants were presented with a percentage indicating the others’ need (high vs. low probability of CPT) before seeing the proposed monetary allocation (need cue, Figure 1B). Thus, in trials with high percentages, the partners had a greater need for money to avoid the painful experience. For each partner, CPT likelihood was low in half of the trials (mean CPT probability of 20%) and high in the other half (mean CPT probability of 80%; random presentation order of high vs. low need trials). As with the monetary proposal amounts, we randomly jittered need-related percentages by 0-4% to minimize habituation and repetition effects.

To ensure the saliency of the experimental need manipulation, all participants completed the painful CPT task themselves before the altruism task outside the scanner. Participants were also told that all three partners had already completed one round of the CPT task before the altruism task and were presented with their ostensible pain ratings (7 on a 7-point scale; 1 = not at all; 7 = extremely). These ratings signaled to participants that all three partners found the ice water equally and extremely painful and were highly motivated to avoid another round of the CPT. After the altruism task, participants completed a variety of computerized sanity check questions and questionnaires outside of the scanner (see Supplemental Figure S2). These sanity checks were used to verify the effectiveness of our experimental merit and need manipulations and to exclude any participants who failed to correctly remember how partners had acted in the behavioral exchange game prior to the altruism task.

## Analysis

### Computational Behavioral Model of Social Perception

We developed two behavioral computational models to characterize individual differences in social perceptions (social perception task) and prosocial behaviors (altruism task). Separately for each task, we modeled participants’ choices and reaction times (RTs) using variants of the drift diffusion model (DDM) ^64^. This model depicts choices as the noisy accumulation of evidence until a sufficient level favoring one choice option is attained. The DDM has been used to examine processes underlying both perceptual and value-based decisions ^64–67^ and is being increasingly applied to studying social and affective decision-making processes ^33, 44, 45, 68–71^.

To model trial-wise responses in the social perception task (Figure 1A), we assumed that when faced with the task of judging each image on a particular dimension (i.e., merit, need, or the control judgments of using both hands), a decision-maker employs the following strategy. At each moment in time, they draw noisy samples of both task-relevant and task-irrelevant evidence about *Merit*, *Need* and *Control* (both hands) from the stimuli, weighted by person- and condition-specific sensitivities *S_merit_*, S*_need_* and *S_control_.* They accumulate these samples of evidence *E* at each timepoint *t* according to the following equation:

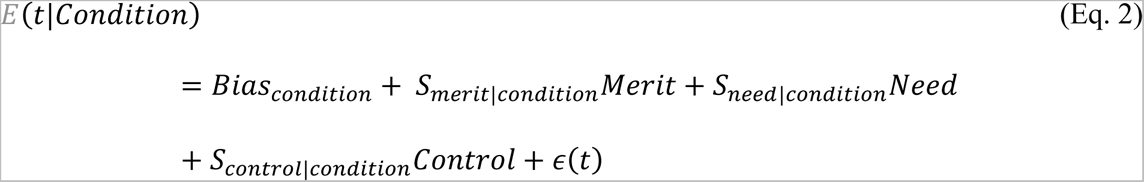

*Condition* refers to the specific judgment (merit, need, control) being performed on that trial (task block). Equation 2 reflects the assumption that a person’s judgment should be most sensitive to task-relevant information (e.g., cues signaling merit during the merit judgment blocks), but might also be inadvertently influenced by task-irrelevant information (e.g., cues signaling need in merit judgment blocks). For example, merit judgments should primarily reflect cues indicating merit (i.e., high sensitivity estimates of *S_merit|merit condition_*) but might also reflect in part cues indicative of need (i.e., a low but non-zero *S_need|merit condition_*). Thus, the model yields a total of nine sensitivity parameters estimated across the three conditions of the social perceptual task. The image-level evidence for *Merit*, *Need* and *Control* (for the stimulus shown on that trial) was estimated using data from an independent participant sample (Supplemental Note S2). For each image, we used the mean-centered average proportion of “yes” responses that the displayed target individual deserved help, needed help, or was using both hands (based on data from the independent sample). We focus on the ‘task-relevant’ estimated parameters *S_merit_* in the merit condition, *S_need_* in the need condition, and *S_control_* in the control condition as indices of participants’ perceptual *sensitivity* to cues suggestive of others’ merit, need, or usage of both hands (but see Supplemental Note S5 for further detail on task-irrelevant sensitivity parameters). Higher parameter values suggest stronger discrimination of the normatively agreed-upon relevant cues (as captured by the independent sample). The DDM also includes three condition-wise free parameters that influence the overall drift, irrespective of the specific image (*Bias_need_*, *Bias_merit_*, and *Bias_control_*; indicated as *Bias_condition_* in Eq. 2). These parameters allow us to capture an individual’s general tendency to identify cues suggestive of others’ merit and need (or control), regardless of the actual social cues present in the image. We will refer to these estimates as perceptual *bias*.

Once the momentary evidence *E* reaches the upper (yes) or lower (no) threshold, evidence accumulation terminates, and the corresponding choice is implemented. The difference between thresholds is estimated by a set of three parameters (*a*_Δ*need*_, *a*_Δ*merit*_, and *a*_Δ*control*_) that represent within-individual stability and difference across tasks (see Supplemental Note S3 for details). The DDM also includes three condition-wise non-decision time (*ndt*) parameters for capturing the time taken to initially encode stimuli and to implement the motor response, estimated similarly to incorporate within-individual stability and change across task conditions. Finally, the model included three condition-wise starting bias (*z*) parameters, which represent another potential mechanism by which biases at the onset of evidence accumulation can impact the decision process. Note that, although we estimated these starting biases to improve model fit, our focus in all primary analyses reported in this paper is on the evidence-related *perceptual* bias for merit and need (represented by *Bias_condition_* in merit and need blocks, respectively), not these motor-related starting biases ^72^. A detailed description of the model fitting procedure is provided in the Supplemental Note S3 and Supplemental Table S1.

### Behavioral Generosity in the Altruism Task

Choices in the altruism task (Figure 1B) involved a trade-off between monetary outcomes for the self and one of three partners. Monetary proposals were selected so that a choice could benefit the partner or the participant, compared to the constant default ($20 for both players). Following previous implementations ^45, 58, 73^, we classified a choice as generous if the participant *accepted* a proposal that benefited the partner at the expense of oneself ($Self < $Other) or *rejected* a proposal that profited themselves at the partner’s expense ($Self > $Other). The overall and condition-specific fraction of generous choices measured participants’ generosity. Differences in generosity across conditions provided model-free estimates of the impact of social cues about others’ merit and need on social behaviors. To assess whether merit and need altered generosity, we computed a mixed-effects logistic regression using the *glmer* function in R. Trial-wise information about generous choice (no = 0, yes = 1) served as the dependent variable. The model included the following fixed effects: a trial-wise indicator for the level of the partners’ need (reference: low need), an indicator for the partners’ merit level (reference: merit low), and their interaction. Participant id was specified as a random effect, as follows:

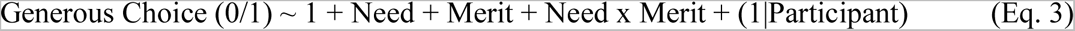

### Computational Behavioral Model of Altruistic Choice

To model choices and reaction time behavior in the 3 (merit: high, unknown, low) x 2 (need: high, low) design of the altruism task, we implemented a second behavioral computational model. Following previous applications ^45, 58^, for each of the six experimental *conditions*, value-related evidence *V* at time *t* was estimated as follows:

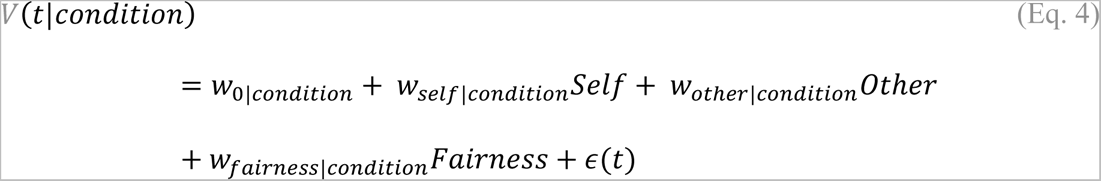

Here, *Self* and *Other* refer to the monetary outcomes of the proposed allocation of a trial (minus the default outcomes of $20 for both; rescaled by dividing by 10). The *Fairness* of the proposed allocation was calculated as −1 × |*Other* − *Self*|. The free parameters for each attribute weight (*w*_*self*_ , *w*_*other*_ , *w*_*fairness*_ ) indicate variance in the degree to which evidence about outcomes to the self, other, or the fairness of the monetary offer guided choices in a particular condition of the altruism task. The value constant w*_0_* represents the extent to which a participant tends to prefer the proposal over the default, regardless of trial-specific values. Matching the computational model of social perception (see above), the model of altruistic choice also estimated free parameters capturing the decision threshold (*a*), non-decision time (*ndt*), and response-related starting bias (*z*) parameter (see Supplemental Note S3 and Supplemental Table S1 for details). The parameters *z*, *ndt,* and *a* were fixed across the merit x need conditions.

To capture variance in weight parameters *w* across the three levels of merit (high, low, unknown) and two levels of need (high, low), we estimated both individual-specific stability and change parameters for each of the four weights *w* (i.e., self, other, fairness, and the value constant). A more complicated model that allowed for need × merit interactions in driving weights did not improve model fit. We thus focus on this simpler model that includes a baseline parameter for each weight, and three change parameters for the effects of increasing or decreasing merit and increasing need (see Supplemental Note S3 for details).

### Correlating Social Perception and Social Action

Next, we tested for a relationship between social perception and social action. To this end, we correlated individual-specific bias and sensitivity parameters obtained from the social perception task with value-based attribute weight parameters in the altruism task. Because we found that some estimates of our computational model diverged from a normal distribution, we used non-parametric statistical tests to examine these relationships. Post-hoc tests were corrected for year> tag found, Please Check. ln:357 cl:32 [Edit].comparisons using the fdr_bh function implemented in MatlabR2022a. All results are reported for 2-tailed statistical tests, unless reported otherwise. We removed outliers from all variables in our analyses based on values that exceeded three standard deviations from the mean.

### fMRI Data

*Acquisition.* All neuroimaging data were acquired at the Caltech Brain Imaging Center using a Siemens Trio 3.0 Tesla scanner outfitted with a 32-channel phased-array head-coil. Functional image acquisition for the social perception task and altruism task occurred on different days in separate sessions (average delay of 303 days). For both fMRI tasks, we acquired gradient echo T2*-weighted echo-planar images (EPIs; 60 slices, voxel resolution 2.5 × 2.5 × 2.5 mm^3^, TR = 700 ms, TE = 30 ms, flip angle = 53°, FOV = 200 mm, interleaved acquisition order, multi-band acceleration factor = 6). We collected 903 volumes for the social perception task and 925 volumes for the altruism task. For all participants, we also acquired a high-resolution anatomical T1-weighted image using a MEMP-RAGE sequence (208 slices, 0.9 × 0.9 × 0.9 mm^3^, matrix size 256 × 256, TR = 2.55 s, TI = 1.15 s, TE = 1.6, 3.5, 5.3, 7.1 ms with RMS echo combination, RAGE flip angle = 8 °). Distortion correction data for the fMRI EPI acquisitions employed a pair of phase-encoding polarity reversed T2w SE-EPI images with identical geometry and EPI echo train timing to the T2*w EPI images (TR 4.8 s, TE 50 ms, flip angle 90°).

#### Preprocessing

Preprocessing of all functional and structural brain data was performed using a standard implementation of fMRIPrep 20.2.3^75^ , which is based on Nipype 1.6.1 ^76, 77^. A detailed description of the standardized analysis steps is provided in Supplemental Note S4.

#### GLM of brain responses in the social perception task

This analysis aimed to identify brain regions recruited during different judgments in the social perception task. For each participant, we estimated a general linear model (GLM) using a canonical hemodynamic response function and a 128s high-pass cut-off filter to eliminate low-frequency drifts in the data. The GLM estimated three regressors of interest corresponding to condition-wise brain responses (implemented in different blocks) during merit inferences (“deserves help?”), need inferences (“needs help?”), and factual control inferences that did not require social inferences (“used both hands?”; Figure 1A). The three condition-wise regressors were defined by the onset time of the first target image and the offset of the final image of each block (two blocks per condition). The GLM also included eight regressors of no interest: six motion regressors, the framewise displacement (estimated during preprocessing of neuroimaging data using fMRIPrep, https://fmriprep.org/), and a session constant. The GLMs were implemented in MATLAB (R2018b) using the SPM12 toolbox (http://www.fil.ion.ucl.ac.uk/spm).

For each participant, we created several contrasts of interest: First, we identified brain regions activated during merit perception (contrast [merit – control]) and need perception (contrast [need – control]). Second, we estimated brain regions that are significantly more (or less) activated for need compared to merit perception (contrasts [merit – need]; [need – merit]). The respective contrast images were then used in four group-level analyses using simple t-tests as implemented in SPM12. Whole-brain maps were thresholded at p < 0.001 (voxel level) and family-wise error (FWE) corrected at a cluster level of p < 0.05. Finally, we tested for brain regions that were reliably activated during both need and merit perceptions, by estimating a formal conjunction of both inference-specific brain maps ([need - control] ∩ [merit - control]) at the group level. Both brain maps used in this conjunction analysis were thresholded at the cluster-forming voxel level of p < 0.001, FWE cluster-level corrected at p < 0.05.

We also estimated a supplemental GLM for fMRI data from the altruism task. Due to COVID-19-related interruptions, only 25 participants from the sample that performed the social perception task also completed the fMRI altruism task. Given the limited sample size and noise level of fMRI data, we decided to focus solely on the behavior in the altruism task to address our research objectives. For completeness, we report the complementary GLM for the fMRI altruism data and its results in Supplemental Note S7.

## Results

This section is structured as follows. First, we report the results of our novel fMRI social perception task to describe how people perceive others’ merit and need. Behaviorally, we used parameter estimates from our computational model of social perception to characterize individuals’ perceptual biases and sensitivities driving variance in social perceptions across people. Neurally, we localized brain regions recruited during merit and need inferences and asked whether neural computations underlying merit and need perceptions are supported by distinct neural circuits or a general-purpose network for social inference processing. Second, we used data from a separate altruism task (collected on average 303 days later) to describe the effects of an interaction partner’s perceived merit and need on meaningful social behavior. Third, we explored if individuals’ perceptual biases and sensitivities (as captured in the social perception task) predicted variance in actual social behavior (observed in the altruism task). We examined this core question about the social perception-action link across time and contexts at both the behavioral and neural levels.

### Social Perception Task: Behavior and Neural Underpinnings of Need and Merit Inferences

Despite viewing the same stimuli, participants differed dramatically in their judgments of others’ merit and need. The percentage of trials perceived by participants as depicting someone as deserving (merit blocks) ranged from 33% to 81%. Similarly, the range for perceiving someone in need (need blocks) was 28% to 73% (Supplemental Figure S3). What mechanisms drive these profound differences in social perceptions?

#### Computational behavioral model of social perception

To address this question, we examined two distinct computational mechanisms: first, people may differ in their general tendency to perceive others as deserving or in need (bias hypothesis). Second, people may differ in their perceptual sensitivity to merit- or need-related evidence in their choice environment and/or their ability to use this information to guide their judgments (sensitivity hypothesis). Importantly, these mechanisms are not mutually exclusive, but may vary in their relative impact on social behaviors across people and contexts. We tested the contribution of these two potential mechanisms using our behavioral computational model of social perception (see Methods).

We first verified that the model adequately accounted for the differences in participants’ choices and response times in all three task conditions (merit, need, and control blocks; for an illustration of the model fit, see Supplemental Figure S4).

Next, we examined the estimated parameters of our hierarchical model at the individual participant level. Positive average sensitivity parameters for merit (*S_merit_* in merit blocks = 2.84 ± 1.02, mean ± std, significantly different from zero, p < 0.001, FDR corrected), need (*S_need_* in need blocks = 3.28 ± 0.75, p < 0.001, FDR corrected), and control (*S_control_* in control blocks = 4.40 ± 0.69, p < 0.001, FDR corrected) verified that participants accurately distinguished between targets who deserve help, need help, and use both hands, and used this perceptual evidence to guide their judgments (Figure 2A). For the estimated perceptual bias parameters, people tended to perceive others as deserving (indicated by an average positive merit bias, *Bias_merit_* = 0.33 ± 0.47, p < 0.001, FDR corrected) and not in need of help (indicated by an average negative need bias, *Bias_need_* = - 0.17 ± 0.39, p = 0.009, FDR corrected). In the control condition, the estimated bias showed a small positive but non-significant trend (*Bias_control_* = 0.09 ± 0.36, p = 0.099, FDR corrected; Figure 2B). It is worth noting that the *Bias* parameters are strongly associated with (though not the sole determinant of) the mean response rate. For a description of the estimated values for the hyper-mean parameters in our model and further sanity checks, see Supplemental Table S2 and Supplemental Note S5.

**Figure 2.**
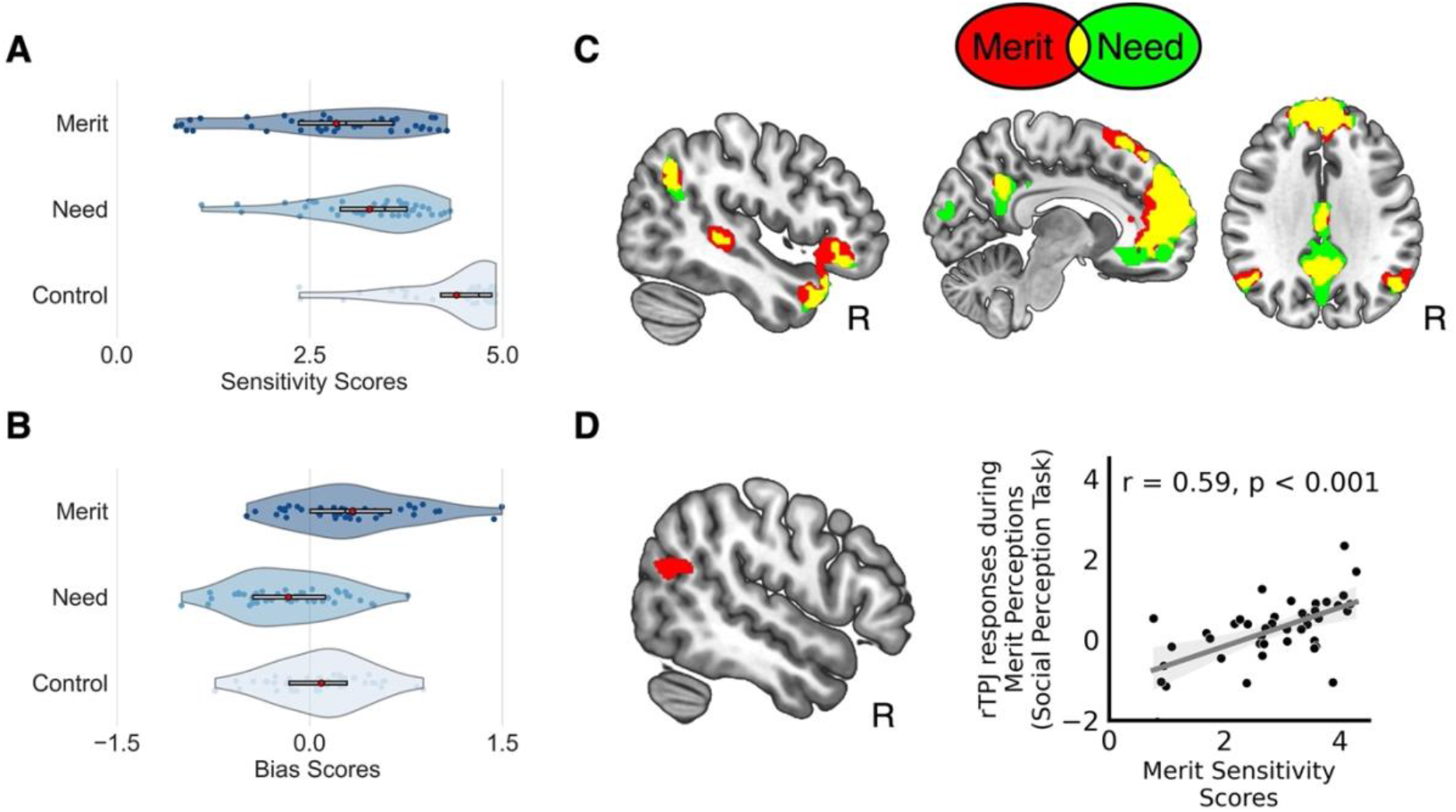
(**A**) Distribution of participant-specific model-based estimates of perceptual sensitivity and (**B**) perceptual bias for each inference condition in the social perception task (merit, need, and control). (**C**) Need and merit inferences activate the mentalizing network to a similar extent (red illustrates brain regions activated for [merit - control], green illustrates brain regions identified for [need - control], and yellow indicates overlap; both contrast maps thresholded at p < 0.001 at the voxel level, FWE corrected at the cluster level at p < 0.05; R = right hemisphere). (**D**) Activity in the right temporoparietal junction (TPJ) during merit perceptions (merit - control) reflects individual differences in merit sensitivity scores estimated in the computational model of social perception.

Notably, individuals’ bias and sensitivity estimates were uncorrelated for merit (Spearman’s r = 0.10, p = 0.73, FDR corrected; *S_merit_* and *Bias_merit_*) and need judgments (Spearman’s r = -0.10, p = 0.73, FDR corrected; *S_need_* and *Bias_need_*), suggesting two distinct mechanisms driving variance in social perceptions across people. Likewise, we found no link between need and merit sensitivity (Spearman’s r = -0.22, p = 0.60, FDR corrected; *S_merit_* and *S_need_*), indicating that people might be sensitive to cues signaling merit but not need, and vice versa. However, estimates of the need bias and merit bias were correlated (Spearman’s r = 0.73, p < 0.001, FDR corrected; *Bias_merit_* and *Bias_need_*), suggesting that people who tend to perceive others as needy might also tend to perceive others as deserving. See Supplemental Figure S5 for full details on intercorrelations of estimates of the computational model. These findings highlight the benefits of formal computational models, which can capture and quantify distinct processes that can be hard to distinguish based on observed behaviors alone. It also raises the interesting question of whether the specificity of need and merit observed at the behavioral level gives rise to inference specificity at the neural level.

#### Neural underpinnings of merit and need perceptions

To address this question, we examined the neural substrates of need and merit inferences obtained in the social perception task. The task is a modified version of an established fMRI why/how task ^53, 54^ that reliably identifies the mentalizing network ^78–82^. Not surprisingly, we found that need and merit inferences also recruited the mentalizing network. The medial prefrontal cortex (MPFC), superior temporal sulcus (STS), temporal pole (TP), temporoparietal junction (TPJ), and posterior cingulate cortex (PCC) were reliably activated during both merit and need inferences, among other regions. Figure 2C illustrates the overlap of brain areas activated during merit and need judgments (Supplemental Table S3 provides the condition-specific results). Table 1 provides the results of the formal conjunction of brain maps identified for [need - control] and [merit - control] inferences, each thresholded at p < 0.001 at the voxel level, FWE corrected at the cluster level at p < 0.05 (see Supplemental Figure S6 for a visualization of the conjunction analysis). No brain region was significantly more activated during merit than need inferences [merit – need] at our omnibus threshold. However, for the reverse contrast [need – merit], need inferences showed significantly greater activation in the cuneus (MNI: [-4, -98, 18], 8741 voxels, t = 7.64), intraparietal sulcus (MNI: [28, -62, 56], 256 voxels, t = 4.58) and sensorimotor cortex (MNI: [4, -38, 64], 294 voxels, t = 4.39) (p < 0.001 at the voxel level, FWE corrected at the cluster level at p < 0.05), suggesting an enhanced activation of the (extended) mirror system ^83, 84^. Future research may reveal additional distinctions between merit and need appraisals in trial-wise (compared to our block-wise) fMRI designs.

**Table 1.** Conjunction of neural activations during social need and merit inferences in the social perception task.

| Brain Region | Side | T | k | MNI |  |  |
| --- | --- | --- | --- | --- | --- | --- |
|  |  |  |  | x | y | z |
| *[Need - Control] ∩ [Merit - Control]* |  |  |  |  |  |  |
| Dorsomedial prefrontal cortex | L | 7.78 | 6250 | -14 | 38 | 48 |
| Superior temporal sulcus | L | 6.61 | 1154 | -64 | -26 | -14 |
| Superior temporal sulcus | R | 5.42 | 146 | 70 | -32 | 0 |
| Temporal pole | R | 6.16 | 523 | 40 | 22 | -32 |
| Posterior cingulate cortex | L | 5.97 | 671 | -2 | -54 | 32 |
| Midcingulate cortex | L | 5.17 | 119 | -2 | -10 | 34 |
| Temporoparietal Junction (TPJ) | R | 4.56 | 167 | 54 | -66 | 34 |
| Temporoparietal Junction (TPJ)/Angular Gyrus | L | 4.58 | 141 | -50 | -62 | 32 |
| Cerebellum | L | 5.33 | 336 | -26 | -82 | -42 |

*Note.* Results are reported at a statistical threshold of $p < 0.001$ at the voxel level, FWE corrected at cluster level at $p < 0.05$ . Only peak activations of each cluster are shown. L = Left hemisphere, R = Right hemisphere, MNI = Montreal Neurological Institute, k = cluster size in voxels.

We also employed supplemental multivariate decoding analyses (searchlight analysis ^85–87^), as commonly used in social perception and neuroscience research ^7, 58, 82, 88, 89^, corroborating our univariate findings (see Supplemental Note S6, Supplemental Table S10). Taken together, our results demonstrate that both need and merit inferences reliably recruited the well-established mentalizing network, and to a comparable extent. Answering our first core question: These findings are consistent with the notion that both appraisals are supported by general-purpose rather than domain-specific social cognitive mechanisms.

#### Model estimates of merit sensitivity modulate the neural underpinnings of merit perceptions

So far, we showed that a general-purpose network involved in social inference processes is recruited during both merit and need inferences. Next, we ran a series of whole-brain analyses to identify brain areas for which inference-evoked brain responses covary with individual differences in estimates of the computational model of social perception. Given the evidence above that individuals activated the social cognitive brain network to make these judgments, we hypothesized that stronger activation in one or more hubs in this network should correlate with greater perceptual sensitivity and/or bias in these social inferences. Consistent with this prediction, we found that brain responses during merit inferences (merit - control) systematically covaried with participants’ merit sensitivity scores in the right temporoparietal junction (rTPJ, peak at [MNI 56, -64, 22], t = 4.24, k = 228 voxels, p < 0.001 at the voxel level, FWE corrected at the cluster level at p < 0.05). In other words, participants with larger merit sensitivity scores (captured in the computational behavioral model) exhibited larger rTPJ responses when appraising someone’s merit (Figure 2D). This effect was specific for brain responses obtained during merit inferences; an analogous whole-brain analysis using neural activation obtained during need inferences (need - control) to predict merit sensitivity did not yield significant effects. We also did not observe any brain regions where inference-evoked brain responses were associated with variance in participants’ need sensitivity or bias scores for either merit or need at the whole brain level (p < 0.001 at the voxel level, FWE corrected at the cluster level at p < 0.05). Thus, despite merit and need inferences both recruiting the general-purpose mentalizing network to an equivalent extent on average, we found some evidence of neuroanatomical specificity for merit inferences when considering estimates of the computational behavioral model (Figure 2D). Notably, this functional link between the rTPJ and merit sensitivity was robust when we repeated the whole-brain analysis for the reduced sample of n = 25 participants with overlapping altruism task data (p < 0.001 at the voxel level, FWE corrected at the cluster level at p < 0.05; for details, see Supplemental Table S4; for illustration, see Supplemental Figure S7).

### Altruism Task Behavior

The results reported above relate to *perceptions* of need and merit; however, they say nothing about how such perceptions might influence decisions to help. In a separate altruism task, we examined how independently manipulating a social target’s merit (high, unknown, low) and need (high, low) alter prosocial behavior. This altruism task was unrelated to the social perception task and was completed on average 303 days later. It allowed us to characterize how people *act* on perceptions of merit and need when deciding whether to give aid to another person.

#### Partner’s need and merit alter generosity in the altruism task

Did the experimental manipulations of another’s need and merit affect people’s generosity during altruistic choice? We addressed this question by fitting a mixed-effects logistic regression model to the observed generous and selfish choices (coded as 1/0) in the altruism task (Eq. 3; see Figure 3A for the average proportion of generous choices across conditions, and Supplemental Table S5 for summary statistics). The model’s total explanatory power was substantial (R^2^ = 0.33) and significantly better than a null model that assumed no effect of need or merit (*χ*^2^ (5, N = 28) = 239.68, p < 0.001). Partners’ merit (*χ*^2^ (2, N = 28) = 45.68, p < 0.001) and need (*χ*^2^ (1, N = 28) = 16.79, p < 0.001) both influenced generosity. On average, subjects were more generous to a partner in high (vs. low) need (beta = 0.40, 95% CI [0.21, 0.58], p < 0.001). They were also more generous to a high (vs. low) merit partner (beta = 0.60, 95% CI [0.42, 0.79], p < 0.001) and an unknown (vs. low) merit partner (beta = 0.52, 95% CI [0.33, 0.71], p < 0.001). No interactions between the level of merit and need were significant (*χ*^2^ (2, N = 28) = 2.77, p = 0.25; need [high] × merit [unknown]: beta = 0.19, 95% CI [-0.06, 0.45], p = 0.142; need [high] × merit [high]: beta = 0.19, 95% CI [-0.07, 0.45], p = 0.147). Likewise, results of a formal model comparison revealed that adding the interaction of need and merit did not improve model fit significantly over a model that only considered the main effects (*χ*^2^ (2, N = 28) = 2.75, p = 0.252; Supplemental Table S6). These findings suggest that need and merit inferences had fully independent effects on social choice. Consistent with this notion, we also found that merit-induced changes in generosity (high-low merit partner) and need-induced changes in generosity (high-low need) were uncorrelated (Spearman r = 0.17, p = 0.377). In other words, people who changed their generous behavior as a function of another’s merit were not necessarily the same as those who changed their behavior in response to another person’s need. Consequently, further analyses focused on the main effects of need and merit on altruistic choice. Overall, the experimental manipulations of another’s need and merit affected people’s generosity during altruistic choice.

**Figure 3.**
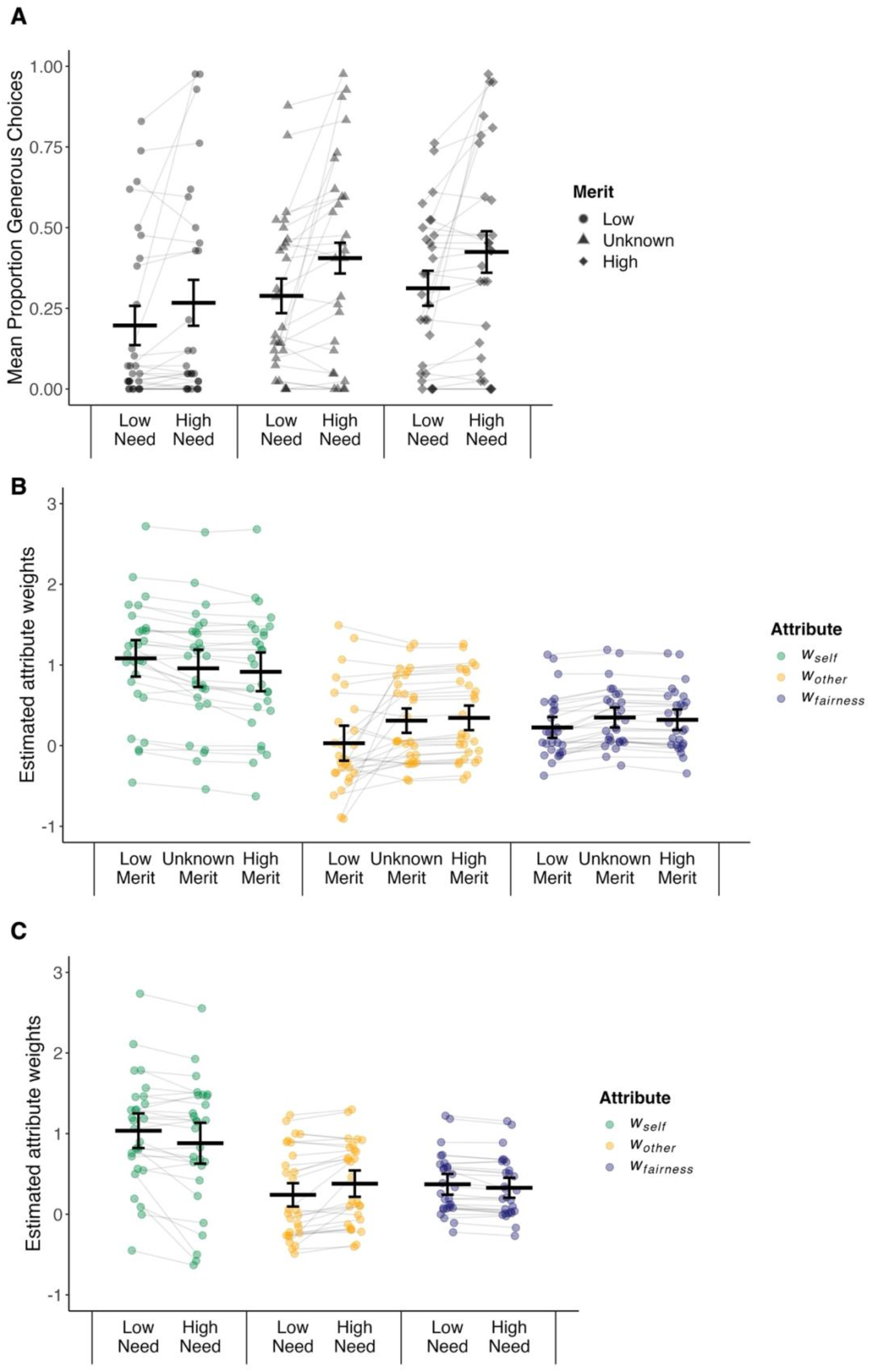
(**A**) Partner’s merit and need altered generosity in the altruism task. High (vs. low) need contexts elicited more generosity (p < 0.001). Compared to a low merit partner (circle), generosity was enhanced towards a high merit (diamond, p < 0.001) and an unknown merit partner (triangle, p < 0.001). (**B**) Condition-specific attribute weights (*w_self_, w_other_, w_fairness_*) for low, unknown, and high merit partner contexts. (**C**) Condition-specific attribute weights for low and high need contexts. All p’s ≤ 0.01, FDR corrected, for the six pair-wise comparisons of changes in attribute weights (high vs. low merit/need). Dots represent participant-specific estimates from the computational model of altruistic choice (see Supplemental Note S3 for detail); black lines illustrate the estimated means and 95% confidence intervals.

**Figure 4.**
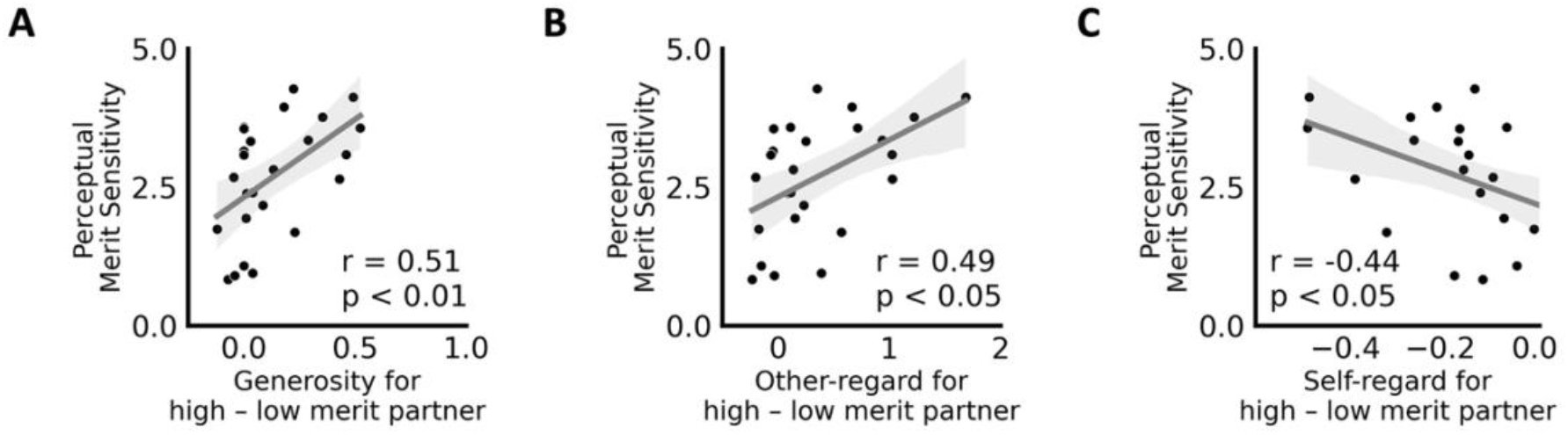
(**A**) Variance in individuals’ general sensitivity to others’ merit (social perception task) is positively linked with merit-related changes in generosity, (**B**) other-regard (*w_other_*), and (**C**) self-regard (*w_self_*) during altruistic choice for high vs. low merit partners. Higher values on the x-axes indicate increased generosity/other-regard and decreased self-interest when interacting with a high vs. low merit partner in the altruism task.

#### Computational behavioral model of altruistic choice: partners’ merit and need alter social attribute weights

Several mechanisms might drive the changes in altruistic choices we observed (see above): others’ merit or need might decrease self-interest, increase other-regard, increase fairness considerations, or some combination of these. To identify the relative contribution of the processes that drive generosity, we turned toward an established behavioral computational model of altruistic choice (Eq. 4; see Methods). We first verified that our computational model of altruistic choice fit the data well by showing that choices and reaction times (RTs) were captured with high accuracy (for visualization of model fit, see Supplemental Figure S4).

Next, we examined the relative importance of the social attributes (drift weights *w*), irrespective of the partner’s need or merit. On average (across all conditions), self-related outcomes strongly guided choices in the altruism task (indicated by the average overall positive weight on payoffs for self, *w_self_* = 0.97 ± 0.75, mean ± std), more so than concern for others’ outcomes (overall *w_other_* = 0.22 ± 0.49, p < 0.001; FDR corrected) or fairness (overall *w_fairness_* = 0.35 ± 0.36, p = 0.028; FDR corrected). These results match the findings of prior studies ^58, 73^. Variance in these *overall* attribute weights across people served as an indicator of more general, context-independent differences in (pro)social decision-making (explaining who will be more generous, on average, and why).

Finally, we tested if contextual cues about the partner’s merit or need altered the degree to which benefits for oneself (*w_self_)*, others (*w_other_*), or fairness concerns (*w_fairness_*) guided social choices (separate Wilcoxon signed rank tests for each attribute and experimental manipulation; FDR corrected for year> tag found, Please Check. ln:357 cl:32 [Edit].comparisons; see Supplemental Table S7 for attribute-specific estimates; see Figure 3B and C for illustration). Any changes in model estimates would indicate *context-dependent* effects (of varying levels of merit and need) of attributes’ input on altruistic behaviors. When interacting with a high (vs. low) merit partner, benefits for oneself (*w_self_*) guided choices *less* (p < 0.001, FDR corrected), whereas considerations of others’ benefits (*w_other_,* p < 0.01, FDR corrected) and fairness concerns (overall *w_fairness_*, p < 0.001, FDR corrected) guided choices *more*. Mirroring this pattern of results, in high (vs. low) need settings, weights on self-related outcomes *decreased* (p < 0.01, FDR corrected), whereas weights on others’ benefits *increased* (p *<* 0.001, FDR corrected). Unlike the merit-induced effects, a partner’s high (vs. low) need *reduced* weights on fairness concerns on choices (p < 0.001, FDR corrected). Thus, if others faced great need, participants were more willing to ignore their fairness preferences. Together, these results suggest that merit- and need-evoked changes in generosity are driven by systematic changes in the social decision process - namely changes in attribute weights - as captured in our behavioral computational model of altruistic choice. Thus, we addressed another key question by showing that partners’ merit and need levels changed how specific choice-relevant considerations (self-regard, other-regard, fairness) guided people’s decisions to act prosocially.

#### Individual differences in effects of others’ need and merit and altruistic decision-making

Notably, people differed substantially in their overall generosity (Supplemental Table S5) and the degree to which generosity varied as a function of their partner’s need and merit. Our model assumes this variation was driven by changes in social attribute weights *w* estimated in the behavioral computational model (see above). To quantify these individual differences and examine their relations, we calculated change scores of model-based estimates for each participant. For example, to capture the change in an individual’s other-regard as a function of their partner’s need, we computed the following participant-specific difference score [*w*_*other*_ in high need – *w*_*other*_ in low need]). Likewise, to assess the change in other-regard in response to their partner’s merit, we estimated the change in [*w*_*other*_ high merit - *w*_*other*_ low merit]. We did this separately for each attribute weight estimated in the computational model of altruistic choices (*w*_*self*_ , *w*_*other*_ , *w*_*fairness*_ ). These change scores reflect the impact of partners’ merit and need on individuals’ altruistic decision process. As a sanity check, we confirmed that changes in attribute weights reflect participant-specific changes in generosity (Supplemental Table S8). In other words, larger (smaller) changes in observed social behaviors can be explained by larger (smaller) changes in estimated attribute weights. These findings provide insight into the precise mechanism by which others’ merit and need effect (pro)social behavior, namely by altering the weighting of certain choice attributes in the decision process. Below, we link these change scores in social behaviors with estimates of social perception (sensitivity, bias).

### Variance in perceptual sensitivity and bias (social perception task) predict variance in prosocial behavior across people and contexts (altruism task)

The previous section contained three main takeaways regarding the separate altruism task. First, partners’ merit and need independently impacted social behaviors (generosity). Second, we identified the mechanism of these context-dependent changes in social behaviors: merit and need altered the relative importance of self- and other-regard, and fairness preferences (captured in attribute weights *w* estimated in our computational model of altruistic choice). Third, we demonstrated that people differ in the degree to which they change their behaviors in response to social cues on others’ merit or need. This raises an important question: what factors determine the impact of others’ need or merit on behavior?

We propose that stable individual differences in social perceptions - as captured in our computational model of social perception and their neural underpinnings - can provide insights into this question. Behaviorally, the computational model decomposes individual differences in social perceptions into bias and sensitivity terms. These model estimates from the social perception task correspond to two mechanisms that drive variation in people’s social perceptions of others’ merit and need. We propose that these stable perceptual mechanisms can, in turn, impact social decision-making. Here, we bring together data from both tasks: the social perception task (which did not require meaningful social behavior towards others) and the altruism task (which included costly social actions). We had two specific hypotheses in mind designing the study: we speculated that merit bias (or need bias) in the social perception task should be related to *average* weights on others’ outcomes in the altruism task. In contrast, estimates of an individual’s sensitivity to merit (or need) should predict the extent to which a person *alters* their weight on others’ outcomes as a function of their partner’s merit (or need). By combining data from both tasks, we explore a fundamental question: do people’s sensitivities and biases during social perceptions translate into subsequent social action? Notably, we see the results below as evidence of *stable* individual differences, since social action (i.e., generosity) was measured on average almost 10 months (∼303 days) after the social perception task.

#### Perceptual merit sensitivity predicts merit-related contextual changes in altruistic choice

Estimates from our computational behavioral model map onto specific hypotheses about the relationship between behaviors observed in both tasks. Perceptual sensitivity estimates reflect individuals’ tendency to sample and integrate evidence about merit (need) during the perceptual decision process. Higher sensitivity estimates will yield higher discriminability between others as a function of their merit (need). Assuming stable individual differences in sensitivity and the impact of categorizing individuals as meritorious or needy on altruistic decision-making, we hypothesized that individuals with higher perceptual merit (need) sensitivity in the social perception task would exhibit a greater change in social behavior depending on the partner’s merit (need) during the altruism task. To test this hypothesis, we correlated individuals’ merit sensitivity scores (social perception task) with merit-induced changes in social behavior (i.e., participant-specific changes in generosity towards [high - low merit] partners in the altruism task). We found that variance in merit sensitivity reflected merit-induced changes in generosity (Spearman’s r = 0.51, p = 0.010; 4A). In other words, individuals generally sensitive to merit information during social perceptions were also more susceptible to merit information during costly social behaviors. Follow-up analyses using estimates of a computational model of the altruism task revealed that this perception-action link was driven by merit-induced changes in other-regard and self-regard during altruistic choice (change in *w*_*other*_ for [high - low merit] partners: Spearman’s r = 0.49, p = 0.045, FDR corrected; change in *w*_*self*_ for [high - low merit] partners: Spearman’s r = -0.44, p = 0.047, FDR corrected; no significant link with merit-related changes in fairness weight, p = 0.784, FDR corrected) (4B and C). In other words, individuals with higher merit sensitivity showed larger discrimination in the value placed on others’ wellbeing (and self-interest) when interacting with supposedly deserving and undeserving people. Next, we repeated this set of analyses for sensitivity scores estimated in the need condition of the social perception task. Here, variance in participants’ need sensitivity did not reflect need-induced changes in generosity (p = 0.745) or need-induced changes in altruistic choice attributes (all p’s > 0.484). However, this absence of effects for need ought to be interpreted with caution, given the comparatively small sample size. Results were qualitatively similar when statistically controlling for the delay between both tasks (partial correlations).

#### Perceptual bias estimates predict individuals’ overall other-regard, self-regard, and fairness considerations in altruistic choice

We also hypothesized that people’s stable perceptual biases in the social perception task might translate into *context-independent* differences in social action (generosity) across people. Put differently, we assumed that individuals who have a *general* tendency to perceive others as deserving/in need (irrespective of social cues present in the environment) should be more willing to help others irrespective of contextual variation in others’ merit/need. We tested this notion by correlating merit and need *bias* parameters with individuals’ overall generosity in the altruism task and the overall weight of choice-relevant attributes (i.e., outcomes for self, *w_self_*; outcome for other, *w_other_*; and fairness, *w_fairness_*). Contrary to our hypothesis, individuals’ merit bias scores were not correlated with overall generosity (p = 0.282). However, we did find that people who tend to perceive others as deserving tended to be more other-oriented overall during altruistic choices: variance in merit bias scores in the social perception task positively correlated with the weights on others’ benefits (*overall w*_*other*_ , Spearman’s r = 0.50, p = 0.035, FDR corrected; marginal positive link with overall fairness concerns in the altruism task, *overall w*_*fairness*_ , Spearman’s r = 0.43, p = 0.053, FDR corrected; no significant link was observed with overall self-regard, p = 0.153, FDR corrected). This finding suggests that stable differences in people’s tendency to perceive others as deserving predicts people’s overall other-regard across different social choice contexts (on average) ten months later. In contrast, variance in individuals’ need bias (social perception task) in our sample did not correlate with overall generosity (p = 0.150) or overall attribute weights (all p’s > 0.076, FDR corrected). As mentioned above, we cannot rule out the possibility that null findings may be due to the comparatively small sample size and should be interpreted cautiously (also see discussion). Results were qualitatively similar when statistically controlling for the delay between both tasks (partial correlations).

#### Neural markers of merit sensitivity predict merit-related behavioral changes during altruistic choice

So far, within the social perception task, we found that merit-evoked neural activation in the rTPJ reflects variance in people’s merit sensitivity (Figure 2D). Moreover, when combining data from both tasks, we showed that individuals’ merit sensitivity (social perception task) predicts merit-related changes in other-regard and self-regard that guide context-dependent changes in social behavior (altruism task) (4B and C). Considering these findings, a post-hoc test examined whether activity in the rTPJ – obtained during merit inferences (social perception task) – also predicts merit-related changes in other-regard (self-regard) in the altruism task (beyond merit sensitivity). We tested this question by using the following equation:

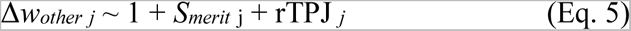

Here, Δ*w_other j_* represents an individual *j’s* merit-related change in other-regard during altruistic choice (*w_other_* for high merit - low merit partners). We used two predictor variables: participants’ behavioral merit-sensitivity scores (*S_merit j_;* estimated in the computational model of social perception) and participant-specific neural responses in the rTPJ obtained during merit inferences (merit - control) in the social perception task (all voxels of the rTPJ cluster, averaged across all voxels in the cluster, Supplemental Table S3). This allowed us to assess the additional predictive power of neural responses in the rTPJ, after controlling for behavioral merit sensitivity. The model’s total explanatory power was substantial (R^2^ = 0.40) and significantly better than a null model with just an intercept (*χ*^2^ (2, N = 25) = 2.46, p < 0.001). We found that both model-based estimates of individuals’ merit sensitivity and rTPJ responses reliably, independently, and to an equivalent degree predict changes in other-regard during social choice approximately 10 months later (*S_merit_*: beta = 0.20, SE = 0.08, 95% CI [0.04, 0.36], t(22) = 2.50, p = 0.021; rTPJ: beta = 0.29, SE = 0.13, 95% CI [0.05, 0.54], t(22) = 2.34, p = 0.029). These findings indicate that neural correlates of merit inferences – namely activity in the rTPJ – predict context-dependent changes in other-regard during social action across time and contexts, above and beyond predictive information related to perceptual merit sensitivity.

For completeness, we also estimated a modified version of Equation 5 in which we changed the dependent variables to merit-evoked changes in self-regard in the altruism task (replacing Δ*w_other j_* with Δ*w_self j_*, defined as participant-specific change scores in self-regard when interacting with a high versus low merit partner). The model yielded an R^2^ of 0.37 and the rTPJ remained a significant predictor (beta = -0.09, 95% CI [-0.17, -0.02], t(21) = -2.47, p = 0.022). However, the behavioral merit sensitivity was reduced to only marginal significance (*S_merit_*: beta = -0.05, 95% CI [-0.10, 0.00], t(21) =-1.92, p = 0.069). Thus, rTPJ responses during merit inferences were tied to estimates of contextual changes in both other- and self-regard to an equivalent or possibly even greater degree than behaviour alone.

## Discussion

Humans do not help others indiscriminately: they are more inclined to help people perceived as needy or deserving ^4, 6–9, 30, 31^. Using a novel fMRI social perception task, we disentangled two distinct computational mechanisms that shape variance in these social judgments: a general *bias* to perceive others as more or less deserving (in need) and a degree of discrimination or *sensitivity* to social cues signaling others’ merit (need). Estimates of these two computations were uncorrelated, suggesting they represent distinct - but not mutually exclusive - processes driving individual differences in perceptions of peoples’ merit and need. We also demonstrated that these computations (for merit, if not need) might be stable and generalizable over time: individuals’ perceptual merit sensitivity predicted the degree to which they discriminated between others based on merit in a separate altruism task completed from 27 to 663 days later. Moreover, their perceptual merit bias predicted a general propensity to weigh others’ outcomes instead of their own during altruistic choices. Neurally, merit sensitivity (but not bias) was associated with increased activity of the TPJ during perceptual judgments, which in turn predicted merit-related discrimination in altruistic behavior. Together, our results identify a set of distinct neurocomputational mechanisms that contribute to our understanding of *when* and *how* perceptions of others translate into social actions.

### Translating perception into action

Variance in people’s sensitivity in merit perceptions predicts context-specific social behaviors and discrimination. This finding contributes to a growing literature regarding parochial altruism. Parochial altruism refers to the tendency to exhibit altruistic behavior towards in-group members and to withhold it from (or even display hostility towards) out-group members ^90^. Parochial altruism occurs around the world in private and public settings ^91^, in sports, politics, war, and religion ^92–95^, and has also been linked to activation patterns in the TPJ ^96–98^. Although we did not use group- or membership-based cues to characterize partners in the altruism task (manipulating merit instead via partner *behaviors* in a separate task), the common locus in the TPJ might indicate shared mechanisms for both types of context-dependent social behaviors. Interestingly, our results suggest a considerable degree of stability of this idiosyncratic perceptual sensitivity across time and contexts, since the social perception and altruism tasks were completed on average ∼10 months apart, up to almost two years for some participants. This finding is consistent with research showing a hereditary component of in-group favoritism and parochial altruism ^99^. Future work should examine the extent to which merit-related perceptual sensitivity represents an innate/genetic or learned quantity, and how it correlates with other types of discriminatory behaviors. Our results also speak to the literature on universal altruism: individual differences in the *bias* to perceive merit correlated with the *overall* weight an individual placed on others during altruistic choice months later. This finding supports empirical evidence that dispositionally cooperative people are more universal in their cooperation ^100^ and suggests that this could partly come from a generalized bias to perceive people as deserving. Although we did not find a distinct neural signature of this bias, future work on genetic or anatomical differences might yield clearer results.

### The value of computational decomposition of perception and action

The power of formal computational models to uncover patterns, principles, and dynamics in social perception and behavior ^101–104^ has made them an increasingly popular tool in economics, psychology, and neuroscience ^105–111^. Our results contribute to this movement by showing how computational models of social perception can provide novel insights into the different computations (i.e., bias and sensitivity) underlying impression formation and its effect on behavior. Although the concept that judgments are a composite of subprocesses is not novel in itself ^112^, modeling allowed us to formally disentangle these different (neuro)computational mechanisms and demonstrate how they shape different aspects of meaningful social action months (or even years) later. Future research should confirm and extend these findings using other social judgments in other, ideally more ecologically relevant contexts ^25^. For example, research suggests that perceptions of others’ warmth and competence can impact hiring decisions ^113, 114^. Are perceptions of warmth or competence likewise driven by stable individual differences in bias and sensitivity? Do stable individual differences in merit bias or sensitivity shape real-world prosociality, such as political support for social welfare programs ^115–117^, or the extent to which different people engage in the online posting of degrading content, harsh comments, or cyber-bullying ^118, 119^?

### The neural bases of need and merit perception

Determining whether socially relevant cues are processed by domain-specific or domain-general neural circuitry remains an active goal of the social and cognitive neurosciences. While considerable evidence exists for specificity in some domains (e.g., face processing ^120–122^ ^122^, emotion recognition ^127–129^, empathy in different modalities ^89^, or even aspects of moral decision-making ^133^), other research points to the broad engagement of the mentalizing network across tasks ^134–140^. Given their importance for decision-making, we sought to determine whether perceptions of either need or merit fall within the category of social phenomena processed by dedicated neural circuits. Our neural findings largely suggest the answer is no. Both merit and need perception engaged the mentalizing network to a similar extent and were virtually indistinguishable neurally, with some minor differences. This held true even when applying multivariate decoding approaches, which have been suggested to be more sensitive than traditional univariate analysis techniques ^141^. We note, however, that perceptions of others’ merit/need in this task likely represent a composite of year> tag found, Please Check. ln:357 cl:32 [Edit].different sub-components (e.g., related to specific social cues sampled and integrated to yield the final social judgment). Thus, our results do not preclude the existence of domain-specific neural circuitry at a lower level of social appraisals (e.g., gender or age categorization, facial and postural cues that a person is in pain, etc.).

### Limitations and future directions

Our results come with some important limitations. One of the biggest concerns is the puzzling absence of any observed neural or behavioral correlates of need sensitivity, in either the social perception or altruism tasks. We primarily relied on univariate analyses to support our conclusions here. Although a supplementary multivariate pattern analysis yielded little additional insight, exploring alternative methods such as the gradient approach or functional connectivity ^142, 143^ could prove more revealing. One other contributing factor to this null result could simply be the small sample size for our altruism task due to COVID-related delays and participant attrition. Other alternative explanations are also possible, however. While purely speculative, one possibility might lie in a limitation of the stimuli used for the social perception task. Need was generally signaled directly and concretely in each picture (i.e., someone with a knife to their throat, someone with a pained facial expression, etc.). In contrast, merit often had to be inferred indirectly from cues about the cognitive and motivational dispositions of the target (e.g., performance of unethical actions, clothing indicating group membership, other signs this person was a “good” person, etc.). The ability to discriminate based on these abstract cues might thus have correlated with TPJ response and with the use of similarly abstract cues during the altruism task to judge merit of social partners. It is possible that if we had signaled need using similarly abstract cues in the social perception task, we might have observed greater associations with mentalizing network regions and/or with altruistic behavior months later. Future work will be needed to more systematically vary factors related to concrete vs. abstract inference, in larger and more diverse samples, and with a greater range of socially relevant behaviors. Finally, another important open question concerns the origins of the identified perceptual biases and sensitivities. Future research should examine if differences in social perceptions stem from societal or religious values see ^144^ conveyed during individuals’ upbringing, genetic factors, or a combination of both.

## Data Availability Statement

Functional imaging data from the social perception task and the altruism task are available at: Conte Social Inference and Context collection (https://nda.nih.gov/edit_collection.html?id=2643). Raw behavioral data for both tasks, ROI masks, computational modeling data used for analysis and to create figures, and the newly created social perception task (including licensing details) are deposited on the Open Science Framework and available at: https://osf.io/4u5vs/.

## Code Availability Statement

FMRI data were preprocessed using the open-source fMRIPrep analysis pipeline (see hyperlinks in the manuscript). Analysis scripts for the first-level GLMs and code for computational models are available on OSF. Source code underlying the supplemental MVPA analysis is openly available here ^145^.

## Acknowledgements

This research was supported by funding from NIMH Conte Center P50 MH094258 (C.A.H., A.T.), the Natural Sciences and Engineering Research Council of Canada RGPIN-2019-04329 (A.T.), and the Social Sciences and Humanities Research Council of Canada 435-2016-1274 (C.A.H.). Computations were performed on the Niagara supercomputer at the SciNet HPC Consortium. SciNet is funded by Innovation, Science and Economic Development Canada; the Digital Research Alliance of Canada; the Ontario Research Fund: Research Excellence; and the University of Toronto. We sincerely thank Julian Michael Tyszka and Tim Armstrong for their support during data collection and Jill Jacobson for helpful comments on the manuscript.

