## Supplementary Material for "A neurocomputational account of the link between social perception and social action"

### Table of Contents

**Supplemental Figures** 4

**Figure S1.** Monetary offers for the altruism task. 4

**Figure S2.** Sanity check of successful merit and need manipulation 5

**Figure S3.** Violin plots of the participant-specific percentage of yes responses for merit, need, and control judgments in the social perception task. 7

**Figure S4.** Model fit of our computational models of social perception and altruistic choice. 8

**Figure S5.** Intercorrelation of estimates of the computational behavioral model of social perception. 9

**Figure S6.** Conjunction of brain areas recruited during need and merit perceptions (social perception task). 10

**Figure S7.** Cluster in the right temporoparietal junction activated during merit perceptions (social perception task) that reflects individual differences in merit sensitivity estimated in the computational model of social perception (in the n = 25 participants with overlapping altruistic choice task data). 11

**Supplemental Tables** 12

**Table S1.** Scaling function and resulting bounded range for each computational parameter in the model for the social perception and altruism task. 12

**Table S2.** Estimates of the computational model of social perception. 13

**Table S3.** Brain regions activated during merit and need judgments in the social perception task. 14

**Table S4.** Brain regions activated during merit perceptions (social perception task) that reflect individual differences in merit sensitivity estimated in the computational model of social perception (in the n = 25 participants with overlapping altruistic choice task data). 15

**Table S5.** Generosity scores across conditions and per condition in the altruism task (n = 28, fraction of trials with generous choices). 16

**Table S6.** Logistic mixed models predicting generous choice. 17

**Table S7.** Model-estimated weights of choice-relevant attributes and drift intercept bias in the altruism task at the participant level (computational model of altruistic choice, n = 28). 18

**Table S8.** Spearman correlations between changes in generosity and changes in parameter estimates (attribute weights) across conditions in the altruism task (n = 28). 19

**Table S9.** Hyper-mean parameter estimates (computational model of altruistic choice). 20

**Table S10.** Whole-brain searchlight decoding of need, merit, and control inferences (social perception task). **21**

**Supplemental Notes** **22**

**Note S1.** Manipulation of partner merit in the altruism task via partner behavior in a separate behavioral task. **22**

**Note S2.** Normative sample description. **24**

**Note S3.** Parameter estimation of the computational behavioral models of social perception and altruistic choice. **25**

**Note S4.** Preprocessing of neuroimaging data. **28**

**Note S5.** Sanity check confirming enhanced perceptual sensitivity scores in task-relevant blocks of the social perception task. **31**

**Note S6.** Multivariate decoding of the inference condition in the social perception task. **32**

**Note S7.** GLM2 for the altruism task and its subsequent analysis. **34**

**References** **36**

**Supplemental Figures**

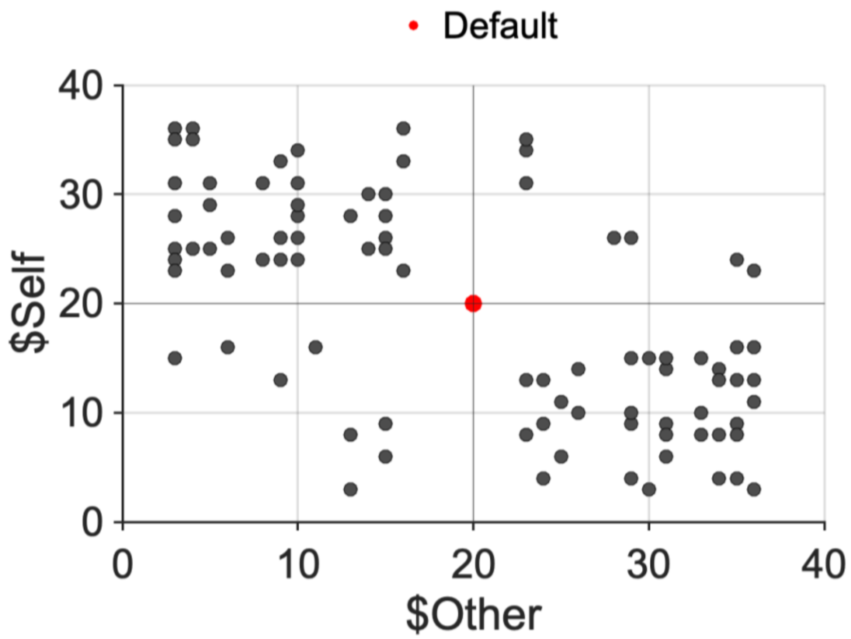


**Figure S1**. Monetary offers for the altruism task (modified dictator game). The monetary pairings of $Self and $Other are indicated as black dots, whereas the default offer ($20 for both) is indicated as a red dot.

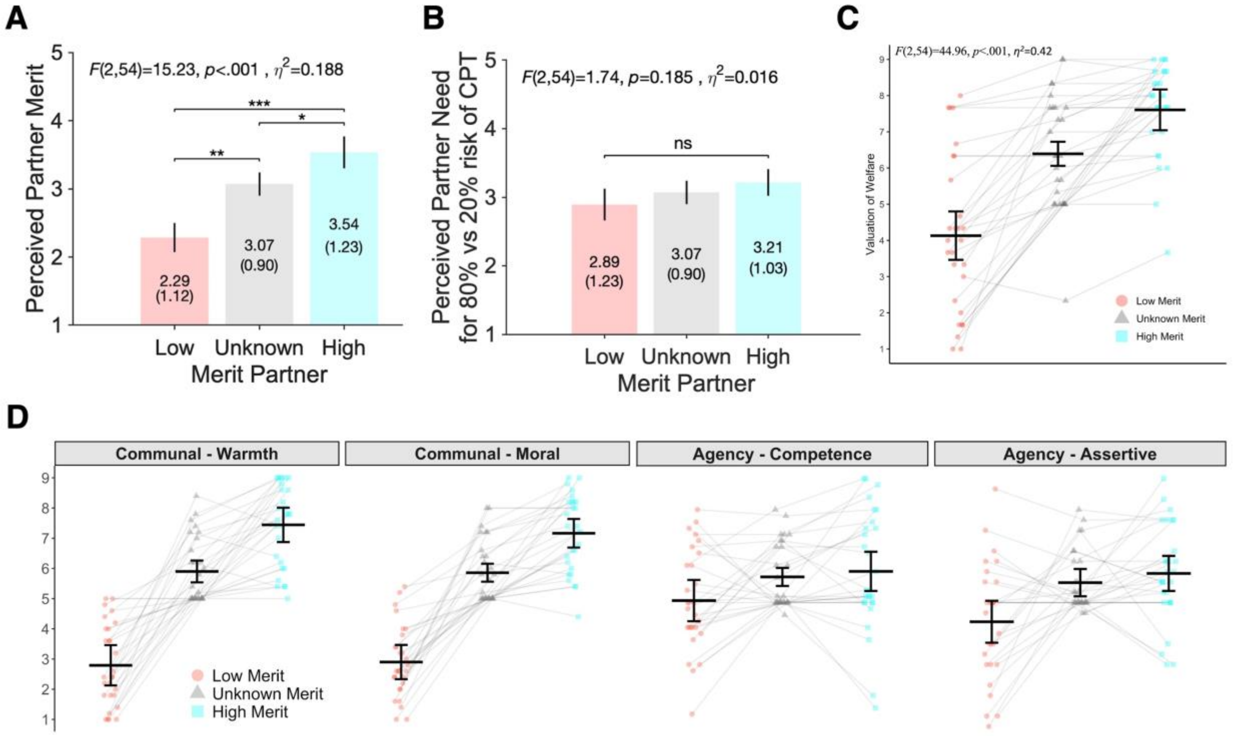


**Figure S2.** Sanity check of successful merit and need manipulation.

We tested for differences in self-reported perceived partner need and merit, obtained at the end of the altruism task (5-point scale, 1 = "Not a t all" to 5 = "Extremely"), using a two-factorial repeated-measures ANOVA. Mauchly's test indicated that the assumption of sphericity had been violated, $W = 0.61$, $p < 0.01$; therefore, the degrees of freedom were corrected using Huynh-Feldt estimates of sphericity ($\varepsilon = 0.75$). (**A)** The results show that perceived partner merit differed significantly across partners, $F(2, 54) = 15.23$, $p < 0.001$, $\eta 2 = 0.188$. Participants perceived a high merit partner as more deserving than an unknown ($p$=0.029) or low ($p$<0.001) merit partner, and a low merit partner was rated as less deserving than an unknown merit partner ($p$=0.003) ("How much more do you think this partner DESERVED to be helped when the chance of having to hold a hand in ice water was 80% versus 20%?"). (**B**) Participants perceived others' need to be greater in high need trials (~80% risk of CPT) compared to low need trials (~20% risk of CPT) by an average of 3.06 (out of a 5-point scale) across all three partners ("How much *more* do you think this partner NEEDED help when the chance of having to hold a hand in ice water was 80% versus 20%"). There were no significant differences in the perceived increased need (for 80% vs 20% CPT) across the three partners ($W = 0.71$, $p < 0.05$, $\varepsilon = 0.81$; $F(2, 54) = 1.74$, $p = 0.185$, $\eta 2 = 0.016$). This was not surprising as participants were told before the altruism task that all three partners perceived the CPT as equally and maximally painful. (**C**) Our experimental merit manipulation also yielded significant differences in self-reported valuation of partners' welfare (Batson et al., 2007), $F(2,54)=44.96$, $p<0.001$, $\eta 2=0.420$ (all $p$'s<0.001, Bonferroni corrected). (**D**) Finally, our merit manipulation affected the perceived trait impressions (Abele et al., 2016) of the three partners in the altruism task (9-point scale, 1="Not at all" to 9="Extremely"). Warmth: $F(2,54)=78.75$, $p<0.001$, $\eta^2=0.707$; Moral: $F(2,54)=95.26$, $p<0.001$, $\eta^2=0.710$; Competence: $F(2,54)=3.41$, $p<0.05$, $\eta^2=0.065$; Assertive: $F(2,54)=8.91$, $p<0.001$, $\eta^2=0.169$. The findings suggest that the merit and need manipulations were effective. Red=Low Merit, Grey=Unknown Merit, Blue=High Merit.

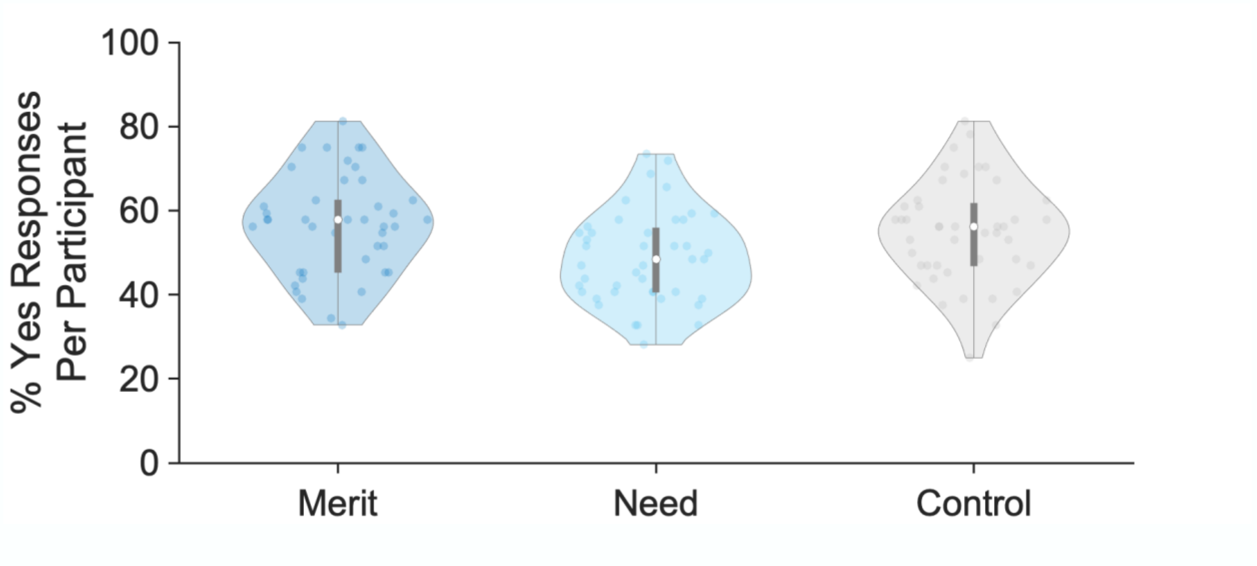


**Figure S3.** Violin plots of the participant-specific percentage of yes responses (dots) for merit (left), need (middle), and control judgments (right) in the social perception task. Merit: 56.70 ± 11.91; Need: 48.69 ± 10.81; Control: 54.87 ± 12.03. Edges of boxplots (grey bars) indicate the 25th–75th percentiles, boxplot whiskers illustrate minima and maxima, and central white dots represent median values. Two outliers for merit and one outlier for need responses were excluded based on values that exceeded three standard deviations from the mean.

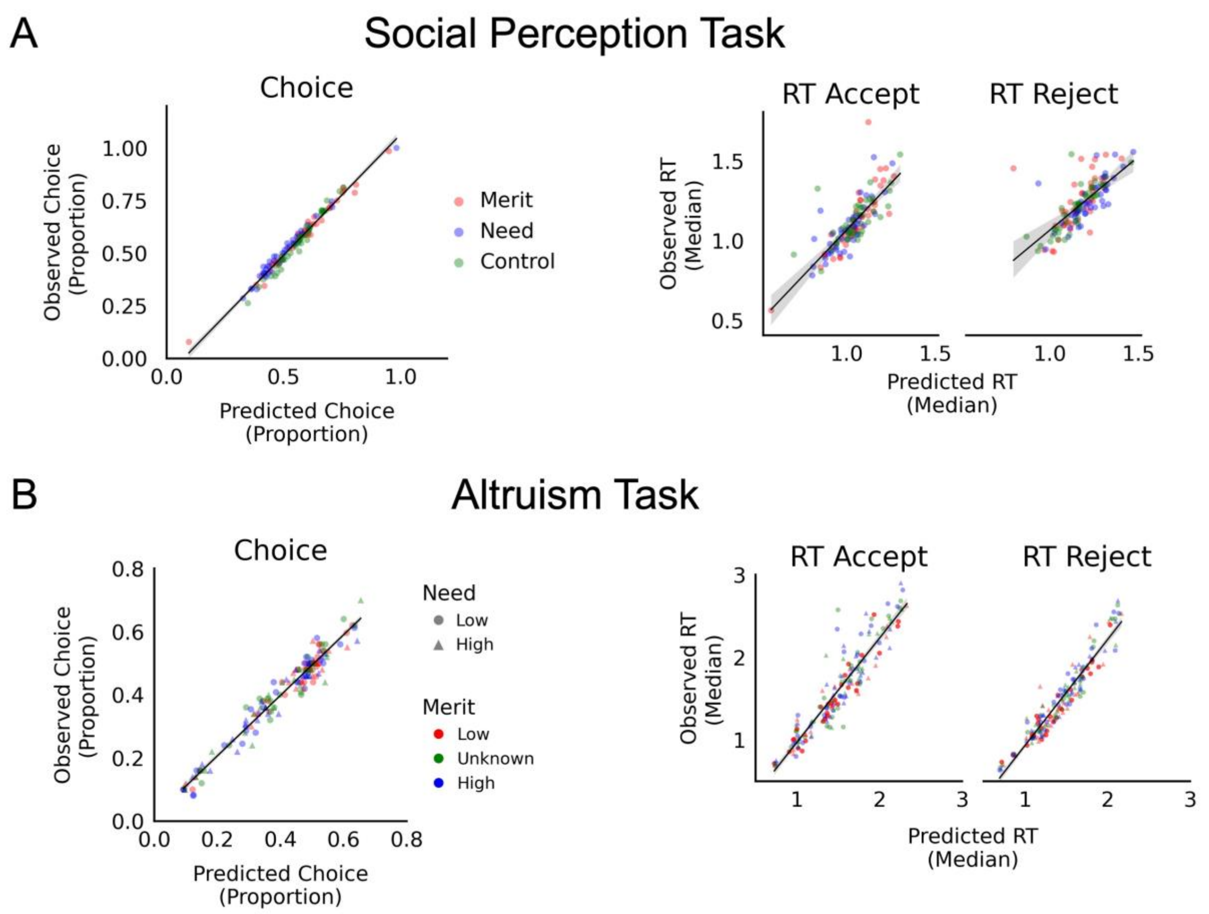


**Figure S4.** Model fit of our computational models of social perception (upper row) and altruistic choice (lower row) illustrated via the correspondence between predicted (model) and observed (data) choices (left panel) and reaction times (RT; middle and right panel).

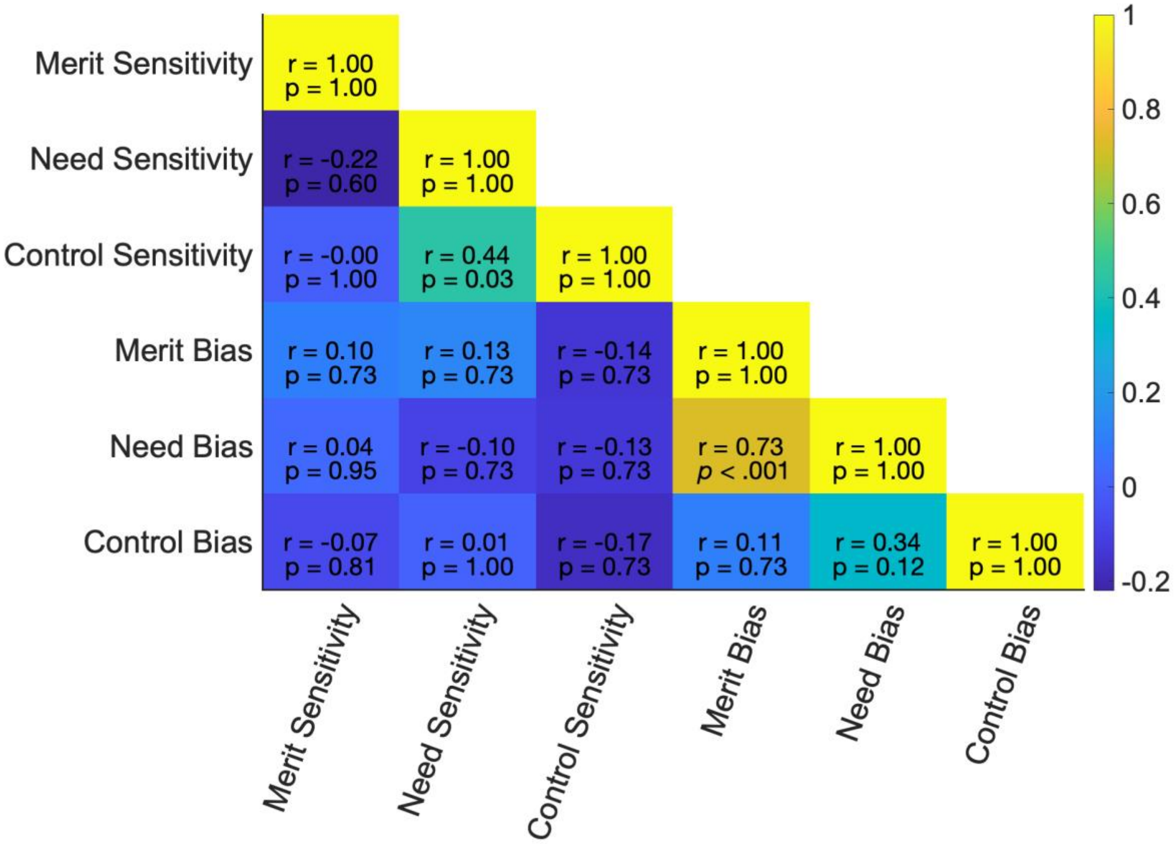


**Figure S5**. Intercorrelation of estimates of the computational behavioral model of social perception (upper values indicate Spearman correlation coefficients, lower values represent p-values, FDR corrected for multiple comparison issues, and pairwise outlier exclusions based on ± 3SD; 1.52% outliers removed for all variables).

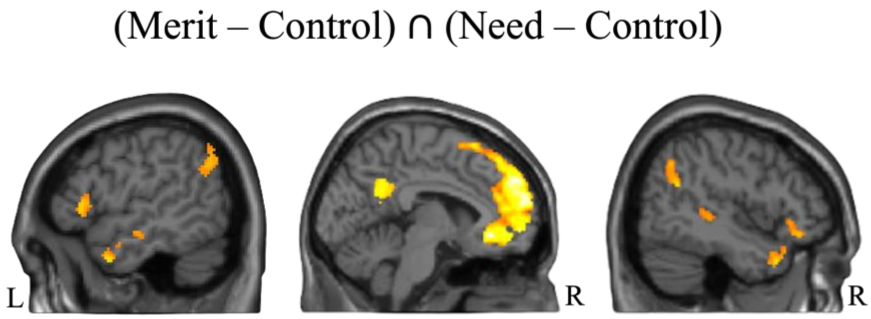


**Figure S6.** Conjunction of brain areas recruited during need and merit perceptions (social perception task) of brain maps identified for [need - control] and [merit - control] inferences (each thresholded at $p < 0.001$ at the voxel level, FWE corrected at the cluster level at $p < 0.05$).

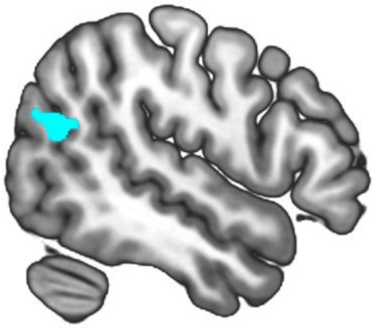

**Figure S7.** Cluster in the right temporoparietal junction (rTPJ) activated during merit perceptions (social perception task) that reflect individual differences in merit sensitivity estimated in the computational model of social perception (in only the n = 25 participants with overlapping altruistic choice task data).

### Supplemental Tables

**Table S1.** Scaling function, $f_{sc}(x)$, and resulting bounded range for each computational parameter in the model for the social perception task (left) and altruism task (right).

| | Social Perception Task | | Altruism Task | |
|---|---|---|---|---|
| **Parameter** | $f_{sc}(x)$ | **Range** | $f_{sc}(x)$ | **Range** |
| all drift parameters ($S$, $w$) | $x \times 10 - 5$ | [-5, 5] | $x \times 10 - 5$ | [-5, 5] |
| $z$ | $x$ | [0, 1] | $x$ | [0, 1] |
| $a$ | $x \times 9.999 - 0.001$ | [0.001, 10.0] | $x \times 9.999 - 0.001$ | [0.001, 10.0] |
| $ndt$ | $x \times 2$ | [0, 2] | $x \times 2$ | [0, 2] |

*Note. S = sensitivity, w* = drift rates, *a* = difference between barriers, *z* = starting bias, *ndt* = non-decision time

**Table S2.** Estimates of the computational model of social perception (hyper-mean parameter estimates, means of the posterior distributions with 95% Highest Density Interval, HDI).

| Parameter | Condition | | |
|---|---|---|---|
| | **Merit** | **Need** | **Control** |
| *bias* | 0.29<br>[0.10, 0.47] | -0.12<br>[-0.31, 0.06] | 0.09<br>[-0.05, 0.23] |
| $S_{merit}$ | **2.94**<br>[2.48, 3.35] | 0.39<br>[-0.07, 0.85] | 0.22<br>[0.11, 0.33] |
| $S_{need}$ | 0.39<br>[-0.002, 0.78] | **3.31**<br>[2.93, 3.64] | -0.12<br>[-0.22, -0.02] |
| $S_{control}$ | 0.42<br>[0.21, 0.64] | 0.26<br>[0.11, 0.41] | **4.57**<br>[4.22, 4.78] |
| *z* | 0.53<br>[0.51, 0.54] | 0.60<br>[0.59, 0.62] | 0.55<br>[0.53, 0.56] |
| *a* | 1.83<br>[1.75, 1.92] | 1.85<br>[1.77, 1.93] | 1.76<br>[1.68, 1.84] |
| *ndt* | 0.59<br>[0.52, 0.66] | 0.62<br>[0.55, 0.69] | 0.60<br>[0.54, 0.67] |

*Note. bias* = overall perceptual bias; $S_{merit}$ = merit sensitivity, $S_{need}$ = need sensitivity, $S_{control}$ = control sensitivity), bold indicates task-relevant sensitivity scores, *z* = starting bias, *a* = difference between barriers, *ndt* = non-decision time

**Table S3.** Brain regions activated during merit and need judgments in the social perception task.

| Brain Region | Side | T | k | MNI (peak) | | |
|---|---|---|---|---|---|---|
| | | | | x | y | z |
| ***[Merit > Control]*** | | | | | | |
| Medial prefrontal cortex | L | 8.04 | 7616 | -8 | 54 | 18 |
| Temporo-parietal junction (TPJ)/ Angular gyrus | R | 4.96 | 278 | 46 | -58 | 34 |
| TPJ/Angular gyrus | L | 4.54 | 267 | -60 | -60 | 34 |
| Superior temporal gyrus | R | 5.84 | 526 | 68 | -34 | -2 |
| Posterior cingulate cortex | L | 6.56 | 687 | -4 | -48 | 28 |
| Medial cingulate cortex | L | 5.29 | 127 | -2 | -14 | 34 |
| Caudate nucleus | L | 5.43 | 74 | -12 | 10 | 12 |
| Anterior insula | L | 7.61 | 1973 | -38 | 20 | -14 |
| Anterior insula | R | 7.35 | 1123 | 34 | 22 | -16 |
| Dorsolateral prefrontal cortex | L | 4.45 | 66 | -34 | 22 | 46 |
| Cerebellum | R | 4.69 | 95 | 26 | -88 | -34 |
| Cerebellum | L | 6.05 | 471 | -26 | -80 | -34 |
| ***[Need > Control]*** | | | | | | |
| Medial prefrontal cortex | L | 7.78 | 7522 | -14 | 38 | 48 |
| TPJ/Angular gyrus | L | 4.58 | 261 | -50 | -62 | 32 |
| TPJ/Angular gyrus | R | 4.56 | 252 | 54 | -66 | 34 |
| Medial temporal cortex | L | 6.61 | 1385 | -64 | -26 | -14 |
| Temporal pole | R | 6.16 | 801 | 40 | 22 | -32 |
| Superior temporal cortex | R | 5.42 | 181 | 70 | -32 | 0 |
| Posterior cingulate cortex | L | 5.97 | 1727 | -2 | -54 | 32 |
| Dorsolateral prefrontal cortex | L | 4.86 | 132 | -40 | 22 | 48 |
| Cerebellum | L | 5.33 | 362 | -26 | -82 | -42 |
| Cerebellum | R | 4.77 | 219 | 26 | -86 | -34 |
| Cuneus | L | 5.34 | 284 | -4 | -100 | 18 |
| ***[Need > Merit]*** | | | | | | |
| Visual cortex | L | 8741 | 7.64 | -4 | -98 | 18 |
| Intraparietal sulcus | R | 256 | 5.24 | 28 | -62 | 56 |
| Somatosensory cortex | R | 294 | 4.97 | 4 | -38 | 64 |

*Note.* Cluster peaks are reported at a statistical threshold of $p < 0.001$ at the voxel level, $p < 0.05$ FWE corrected at the cluster level. There were no significant results for [Merit – Control] > [Need – Control]. L = Left hemisphere, R = Right hemisphere, MNI = Montreal Neurological Institute, k = cluster size in voxels.

**Table S4.** Brain regions activated during merit perceptions (social perception task) that reflect individual differences in merit sensitivity ($S_{merit}$) estimated in the computational model of social perception (in the n = 25 participants with overlapping altruistic choice task data).

| | Brain Region | Side | T | k | MNI (peak) | | |
|---|---|---|---|---|---|---|---|
| | | | | | x | y | z |
| ***Merit Sensitivity*** | | | | | | | |
| [Merit > Control] | Temporoparietal junction | R | 4.48 | 157 | 48 | -62 | 22 |
| | Precuneus | R | 5.66 | 380 | 6 | -60 | 56 |
| | Frontal eye fields | R | 5.12 | 342 | 22 | 0 | 54 |
| | Premotor cortex | R | 5.84 | 113 | 34 | -46 | 58 |
| [Need > Control] | - | - | - | - | - | - | - |
| [Merit > Need] | - | - | - | - | - | - | - |

*Note.* Cluster peaks are reported at a statistical threshold of $p < 0.001$ at the voxel level, $p < 0.05$ FWE corrected at the cluster level. L=Left hemisphere, R=Right hemisphere, MNI=Montreal Neurological Institute, k=cluster size in voxels.

**Table S5.** Generosity scores across conditions (overall) and per condition in the altruism task (n = 28, fraction of trials with generous choices).

| Condition | Mean | SD | Min | Max |
|---|---|---|---|---|
| Overall generosity | 0.34 | 0.23 | 0.08 | 0.91 |
| merit low, need low | 0.24 | 0.24 | 0.06 | 0.86 |
| merit low, need high | 0.30 | 0.30 | 0.00 | 0.98 |
| merit unknown, need low | 0.32 | 0.22 | 0.08 | 0.90 |
| merit unknown, need high | 0.43 | 0.28 | 0.06 | 0.94 |
| merit high, need low | 0.34 | 0.21 | 0.06 | 0.80 |
| merit high, need high | 0.44 | 0.29 | 0.08 | 0.96 |

**Table S6.** Logistic mixed models predicting generous choice (0/1).

| | Model 1 (Null Model) | | Model 2 (Winning Model) | | Model 3 (Interaction Model) | |
|---|---|---|---|---|---|---|
| | Estimate | S.E. | Estimate | S.E. | Estimate | S.E. |
| (Intercept) | **-0.81*** [-1.25, -0.37]** | 0.23 | **-1.55*** [-2.01, -1.08]** | 0.24 | **-1.48*** [-1.95, -1.00]** | 0.24 |
| need [high] | - | - | **0.53*** [0.43, 0.63]** | 0.05 | **0.40*** [0.21, 0.58]** | 0.10 |
| merit [unknown] | - | - | **0.63*** [0.50, 0.75]** | 0.07 | **0.52*** [0.33, 0.71]** | 0.10 |
| merit [high] | - | - | **0.71*** [0.58, 0.83]** | 0.07 | **0.60*** [0.42, 0.79]** | 0.10 |
| need [high] × merit [unknown] | - | - | - | - | 0.19 [-0.06, 0.45] | 0.13 |
| need [high] × merit [high] | - | - | - | - | 0.19 [-0.07, 0.45] | 0.13 |
| self | - | - | - | - | - | - |
| other | - | - | - | - | - | - |
| fairness | - | - | - | - | - | - |
| $R^2$ | 0 | | 0.33 | | 0.33 | |
| AIC | 8973 | | 8742 | | 8743 | |
| BIC | 8987 | | 8777 | | 8792 | |

*Note.* Significant effects indicated in bold. * p < 0.05 ** p < 0.01 *** p < 0.001. Sample size n = 28. The model's intercept corresponds to need = low, merit = low, self = 0, other = 0 and fairness = 0. 95% Confidence Intervals (displayed in brackets) and p-values were computed using a Wald z-distribution approximation.

**Table S7**. Model-estimated weights of choice-relevant attributes ($w_{self}$, $w_{other}$, $w_{fairness}$) and drift intercept bias ($w_0$) in the altruism task at the participant level (computational model of altruistic choice, n = 28).

| Attributes | Estimates | Mean | SD |
|---|---|---|---|
| $w_0$ | Baseline | 0.34 | 0.44 |
| | $\Delta_{Need}$ | -0.03 | 0.06 |
| | $\Delta_{High\ Merit}$ | -0.09 | 0.02 |
| | $\Delta_{Low\ Merit}$ | -0.12 | 0.04 |
| $w_{self}$ | Baseline | 0.96 | 0.72 |
| | $\Delta_{Need}$ | -0.08 | 0.13 |
| | $\Delta_{High\ Merit}$ | -0.04 | 0.07 |
| | $\Delta_{Low\ Merit}$ | 0.12 | 0.13 |
| $w_{other}$ | Baseline | 0.31 | 0.52 |
| | $\Delta_{Need}$ | 0.07 | 0.10 |
| | $\Delta_{High\ Merit}$ | 0.03 | 0.03 |
| | $\Delta_{Low\ Merit}$ | -0.28 | 0.48 |
| $w_{fairness}$ | Baseline | 0.35 | 0.37 |
| | $\Delta_{Need}$ | -0.02 | 0.03 |
| | $\Delta_{High\ Merit}$ | -0.03 | 0.03 |
| | $\Delta_{Low\ Merit}$ | -0.12 | 0.10 |

*Note*. For hyper-mean parameter estimates of the computational model of altruistic choice (means of the posterior distributions with 95% Highest Density Interval, HDI), see Supplemental Table S9. To reconstruct Figure 3B, take [Baseline + $\Delta_{Low\ Merit}$], [Baseline], and [Baseline + $\Delta_{High\ Merit}$]. To reconstruct Figure 3C, take [Baseline – $\Delta_{Need}$], and [Baseline + $\Delta_{Need}$].

**Table S8.** Spearman correlations between changes in generosity and changes in parameter estimates of the computational model of altruistic choice (attribute weights) across conditions in the altruism task (FDR corrected, n = 28).

| | | Merit-induced Changes in Generosity [High Merit – Low Merit] | | Need-induced Changes in Generosity [High Need - Low Need] | |
|---|---|---|---|---|---|
| Condition Altruism Task | Changes in Parameter Estimates | R coefficient | P-value | R coefficient | P-value |
| Merit [High-Low] | $\Delta w_{self}$ | **-0.85** | **0.000 *** | -0.19 | 0. 344 |
| | $\Delta w_{other}$ | **0.91** | **0.000 *** | 0.01 | 0.948 |
| | $\Delta w_{fairness}$ | **0.56** | **0.018*** | -0.11 | 0.569 |
| Need [High-Low] | $\Delta w_{self}$ | 0.19 | 0.333 | **0.75** | **0.000 *** |
| | $\Delta w_{other}$ | -0.14 | 0.484 | **-0.88** | **0.000 *** |
| | $\Delta w_{fairness}$ | -0.11 | 0.570 | -0.03 | 0.895 |

*Note*. Individuals' merit-induced changes in generosity were reflected by individuals' altered weights for self-interest (payoffs for self), other-regard (payoffs for partners) and fairness considerations on choices. Need-related changes in generosity were reflected by altered consideration of benefits for self and others (but not fairness). In other words, individuals who were most sensitive to information about others' merit and need (i.e., large differences in generosity towards a high vs. low merit/need partner) changed their decision process more strongly (captured in larger shifts in the weights on outcomes for self, other, and fairness on choices as a function of others' merit/need). These findings provide insights into the precise mechanism by which need and merit affect (pro)social behavior, namely by altering the weighting of certain choice attributes in the decision-making process.

**Table S9.** Hyper-mean parameter estimates (computational model of altruistic choice).

| Parameter | Fixed | Low Need | High Need | Low Merit | High Merit |
|---|---|---|---|---|---|
| $w_0$ | – | 0.38<br>[0.16, 0.59] | 0.31<br>[0.09, 0.52] | -0.12<br>[-0.19, -0.06] | -0.10<br>[-0.14, -0.05] |
| $w_{self}$ | – | 1.05<br>[0.74, 1.35] | 0.89<br>[0.58, 1.20] | 0.12<br>[0.02, 0.22] | -0.04<br>[-0.11, 0.03] |
| $w_{other}$ | – | 0.24<br>[0.02, 0.47] | 0.38<br>[0.16, 0.61] | -0.28<br>[-0.49, -0.07] | 0.03<br>[-0.01, 0.08] |
| $w_{fairness}$ | – | 0.37<br>[0.21, 0.53] | 0.33<br>[0.17, 0.49] | -0.12<br>[-0.18, -0.06] | -0.03<br>[-0.06, 0.00] |
| $z$ | 0.50<br>[0.48, 0.53] | – | – | – | – |
| $a$ | 2.61<br>[2.43, 2.80] | – | – | – | – |
| $ndt$ | 0.61<br>[0.53, 0.69] | – | – | – | – |

*Note.* Means of the posterior distributions with 95% HDI in brackets. $w$ = drift rates ($w_0$ = bias, $w_{self}$ = self outcome, $w_{other}$ = partner outcome, $w_{fairness}$ = fairness between self and partner outcome), $z$ = starting bias, $a$ = difference between barriers, $ndt$ = non-decision time. To capture how attribute weights ($w_{self}$, $w_{other}$, $w_{fairness}$) differed across conditions, our computational model estimated four separate drift parameters for each attribute: 1) baseline sensitivity for the unknown partner; 2) an additive term related to the two levels of need (high = +1, low = -1); 3) an additive term indicating high-merit partner trials (coded as 1/0); and 4) an additive term indicating low-merit partner trials (coded as 1/0). The $z$, $a$, and $ndt$ parameters were fixed across need and merit conditions.

**Table S10.** Whole-brain searchlight decoding of need, merit, and control inferences (social perception task).

| | Brain Region | Side | T | k | MNI (peak) | | |
|---|---|---|---|---|---|---|---|
| | | | | | x | y | z |
| [Merit vs. Control] | | | | | | | |
| | TPJ (temporoparietal junction) / angular gyrus | L | 6.08 | 3885 | -52 | -64 | 44 |
| | TPJ / angular gyrus | R | 5.48 | 2241 | 48 | -60 | 32 |
| | DMPFC (dorsomedial prefrontal cortex) | L | 5.37 | 1129 | -6 | 68 | 12 |
| | DMPFC | R | 4.94 | 1265 | 14 | 42 | 40 |
| | Superior frontal sulcus | R | 5.05 | 678 | 26 | 12 | 60 |
| [Need vs. Control] | | | | | | | |
| | Occipito-temporal cortex *Note: the cluster includes the bilateral TPJ, DMPFC, temporal poles and posterior cingulate cortex | L/R | 10.94 | 129.326 | -56 | -60 | -10 |
| [Merit vs. Need] | | | | | | | |
| | - | - | - | - | - | - | - |

*Note.* Results are reported at a statistical threshold of $p < 0.001$ at the voxel level, FWE corrected at the cluster level at $p < 0.05$. L = Left hemisphere, R = Right hemisphere, MNI = Montreal Neurological Institute, k = cluster size in voxels. Only peak coordinates of clusters are reported. Sample size n = 44.

**Supplemental Notes**

**Note S1.** Manipulation of partner merit in the altruism task via partner behavior in a separate behavioral task.

Before the altruism task (outside of the scanner), all participants completed a separate behavioral exchange game (sequential iterated Prisoner's Dilemma, modified from (Singer et al., 2004; Singer et al., 2006)). We used this separate task to experimentally manipulate partner merit for the subsequent altruism task. Participants were first informed about the game's overall structure, as follows: two anonymous players each start with 10 points. One player is randomly selected to act first and chooses how many points to send to the other player. Each sent point is tripled (e.g., if the player sends 10 points, the other player receives 30). The second player is then informed of the first player's choice and chooses how many of their total points to send in return (again, anything sent is tripled). Total joint earnings are maximized if each player sends the maximum number of points to the other player. Importantly, the second player can decide not to send any points back at the expense of the first player. Participants were informed that they would be paired with a different person (ostensibly another participant) for each trial. After the instruction, all participants were "randomly" assigned to act as the second player. Participants completed ten trials where each first player sent one of the following amounts: 1, 3, 5, 5, 7, 7, 9, 9, 10, and 10. The ordering of these trials was randomized across participants.

We implemented the following approach to manipulate the perceived merit of three anonymous players. After playing the game themselves, participants received feedback about the choices of two other players and had to guess the choice of a third player. Participants were informed that these three individuals had played the sequential prisoners' dilemma at the same time as the participant but in different rooms. Participants were informed that player 1 behaved uncooperatively ("Sent 0 out of 10, which tripled to become 0"; indicating low merit), and player 2 behaved cooperatively ("Sent 10 out of 10, which tripled to become 30"; indicating high merit). As a control condition, no information about player 3 was provided (unknown merit); instead, participants were asked to guess that player's choices.

Subsequently, participants completed self-reported ratings on a scale from 1 ("Not at all") to 9 ("Extremely") regarding perceived partner traits (morality, warmth, competence, assertiveness) and valuation of partner welfare. These ratings indicated that the merit manipulation was successful (Supplemental Figure S2C and D). Participants were then informed that they would play the next task (altruism task) in the scanner with those three partners, represented by three colored geometric shapes.

To increase believability that participants were indeed interacting with three other partners, when they arrived at the laboratory, the experimenter guided them to the room where the prescanner tasks were completed. As the participants were led to the room, they and the experimenter passed another room with the door ajar and a recording playing from within of an ostensible conversation between another experimenter and participant. The real experimenter drew participants' attention to this as they passed and mentioned that the other participants had already arrived.

**Note S2.** Normative sample description.

Estimating participants' sensitivity to need and merit cues in the social perception task requires quantifying the degree of need and merit displayed in each picture. We assessed this quantity by obtaining normative ratings from an independent sample of participants recruited through Mechanical Turk (MTurk; http://www.mturk.com) and Qualtrics (http://www.qualtrics.com). The normative sample included 50 participants (17 females; mean age = 42 years, range = 25–67; 82% of White race; 100% native English speakers). This sample performed a behavioral online version of the social perception task. Participants made binary judgments of whether the displayed target individual needed help, deserved help, or used both hands (control) (i.e., the same judgment made by our main fMRI sample). For a given judgment (need/merit/control), the mean proportions of 'yes' responses across the sample for each image were used to operationalize perceptual evidence on merit, need, and control in the experimental stimuli used in the social perception task. We used these data from the separate sample to estimate the free sensitivity parameters in the behavioral computational model of social perception described in the main text. Data from our independent normative participant sample is available on the Open Science Framework (OSF; see https://osf.io/4u5vs/).

**Note S3.** Parameter estimation of the computational behavioral models of social perception and altruistic choice.

For both the social perception task and the altruism task, we identified the best-fitting parameter values by estimating their posterior distributions with hierarchical Bayesian models using differential evolution Markov chain Monte Carlo (DE-MCMC) sampling (Holmes & Trueblood, 2018; Turner et al., 2013). Specifically, we used an analytic solution (Navarro & Fuss, 2009) to calculate the likelihood of the observed data (i.e., choices and RTs) given a combination of parameter values and used this likelihood to construct a Bayesian estimate of the posterior distribution of the likelihood of the parameter values given the data. To maximize the amount of data used for fitting our model, we included trials in which participants did not respond before response deadlines by estimating the probability that simulations would fail to generate a choice within this time frame. Before model fitting, we removed a few trials where participants made a response in less than 200 ms (0% and 0.02% of trials in the social perception task and altruism task, respectively).

At the participant-level of our hierarchical model for the *social perception task*, we fit 7 parameters [$bias$, $S_{need}$, $S_{merit}$, $S_{control}$, $z$, $a$, $ndt$] for each of the task conditions (need/merit/control judgments) simultaneously. Specifically, the drift sensitivity parameters were estimated separately for each of the three task conditions (merit, need, control) and, thus, their values were independent across conditions. However, because we expected a participant's thresholds, starting biases, and non-decision times to be correlated across conditions, we used a method involving a "baseline and difference" approach, in which one parameter specified the value obtained in the baseline/control (hands) condition, and two additional parameter governed the *difference* between the control value for that parameter and the specific task condition (i.e., merit – baseline, need – baseline). Thus, we obtained 21 parameter values per participant (7 free parameters x 3 conditions). At the population-level of our model, we obtained estimates of the hyper means and hyper standard deviations for each of these parameters, resulting in 42 hyper parameters.

At the participant-level of our hierarchical model for the *altruism task*, we fit 3 parameters [$z$, $a$, $ndt$] that were fixed across the need × merit conditions. In addition, we used a combination of 4 parameters [$w_0$, $w_{self}$, $w_{other}$, $w_{fairness}$] that were allowed to vary across conditions to describe the value-based evidence accumulation (drift) term in the model. For these parameters, we estimate only the independent main effects for need and merit, since another model that allowed for need × merit interactions did not improve model fit. Although the four weight parameters [$w_0$, $w_{self}$, $w_{other}$, $w_{fairness}$] were allowed to vary across the 6 combinations of need and merit, we also assumed that each of these parameters might be related within a single participant. Thus, we again took a "baseline and difference" approach to capture how weight parameters differed across conditions. More specifically, our DDM model estimated 4 separate drift parameters for each attribute: *(i)* baseline sensitivity for the unknown partner; *(ii)* an additive term related to difference in the need level ($\Delta_{Need}$, high = +1, low = -1); *(iii)* an additive term indicating high-merit partner trials, coded as 1/0; and *(iv)* an additive term indicating low-merit partner trials, coded as 1/0. For ease of comparison, when reporting the estimates for the main effect of need, we recode parameter estimates for high need as (baseline + $\Delta_{\text{Need}}$, see *i* and *ii* above) and low need as (baseline - $\Delta_{\text{Need}}$, see *i* and *ii* above). Thus, in total, we obtained 19 parameter values per participant. At the population-level of our model we obtained estimates of the hyper means and hyper standard deviations for each of these parameters, resulting in 38 hyper parameters.

In total, we ran 3 × *k* chains in parallel for each model, where *k* is the number of participant-level parameters per participant (i.e., social perception task: 63 chains; altruism task: 57 chains; Holmes & Trueblood, 2018). To preserve within-individual consistency for select parameter values (see above), we fit all conditions simultaneously. Additionally, we restricted possible parameter values as shown in Supplemental Table S1. Given these constraints, we employed a transformation in parameter sampling to ensure that the prior distributions of the hyper mean model parameters were truly uniform (noninformative) across the specified range in each condition.

Specifically, for the social perception task, the DE-MCMC sampler sampled 3 MCMC model parameters each for *ndt*, *a*, and *z* – one for each condition (i.e., need, merit, and control). To preserve within-participant consistency across the three conditions while ensuring that an inverse probit transformation would result in a uniform distribution across values of [0,1], we computed the difference between one MCMC parameter and the sum of the other two (e.g., $ndt_{need}$ – ($ndt_{merit}$ + $ndt_{control}$)) and then transformed this value via an inverse probit transformation. In this way, by specifying the priors of the MCMC hyper mean model parameters as normal distributions with mean of 0 and variance of 1/3, *N*(0, 1/3), their combination and subsequent inverse probit transformation yielded a uniform distribution across values of [0, 1] for parameters in each condition. These transformed values were then scaled by their respective functions, $f_{sc}(x)$, to the range of values as seen in Supplemental Table S1 to derive the model parameters. The priors of the MCMC hyper standard deviation parameters were specified as gamma distributions $\boldsymbol{\Gamma}$[1, 1].

For the altruism task, the DE-MCMC sampler sampled MCMC model drift weight (*w*) parameters composed of stability ($M_{\text{params}}$) and effect of need and merit manipulations (change across conditions; $\delta_{\text{params}}$). In the case of need-related effects, the sum of these MCMC model parameters $M_{\text{params}} + \delta_{\text{params}} \times$ Condition (effects coded as high need = 1; low need = -1) is then transformed via an inverse probit transformation. The priors of the MCMC hyper mean model parameters for both $M_{\text{params}}$ and $\delta_{\text{params}}$ were specified as normal distributions with mean of 0 and variance of 0.5, *N*(0, 0.5), because inverse probit transformations of the sum of two normal distributions with mean 0 and variance 0.5 yields a uniform distribution across values of [0, 1] for parameters in both conditions. These transformed values were then scaled by their respective functions, $f_{sc}(x)$, to the range of values as seen in Supplemental Table S1 to derive the diffusion model parameters. For the high and low merit conditions, the corresponding drift weight parameter was added following probit transformation. The priors of the MCMC hyper standard deviation parameters were specified as gamma distributions $\boldsymbol{\Gamma}[1, 1]$.

To construct the estimated posterior distributions of each parameter, we sampled 5,000 iterations after an initial burn-in period (burn-in of 15,000 for the social perception task, 10,000 for the altruism task). We then thinned these 5,000 samples by only keeping every 5th sample, leaving 1,000 samples per chain. For each iteration, the DE-MCMC algorithm proposes a new set of parameter values for each chain based on the scaled difference between two other randomly selected chains (Turner et al., 2013). The scaling for the difference between parameter values was $\gamma = 2.38/\sqrt{2d}$ where $d$ is the number of parameters in the parameter space (e.g., for the social perception task, 42 at the population-level and 21 at the participant-level; Turner et al., 2013). At the hyper-level, parameter proposals were blocked such that one new parameter value was proposed at a time. At the participant-level, parameter proposals were unblocked (Turner et al., 2013). The newly proposed parameter values are then evaluated by the Metropolis-Hastings algorithm for inclusion in the posterior distribution. Additionally, we implemented a probabilistic migration step, $\alpha = 0.01$, at each MCMC step instead of the differential evolution to improve chain-mixing and convergence towards the high probability density region of the posterior distribution of parameters. The migration step cycles the positions of a subset of chains ($N_{\text{migrate}}$ uniformly sampled from the total number of chains) such that the positions of chains were compared against $\{i + 1, i + 2, \ldots j, i\}$ and evaluated based on the Metropolis-Hastings algorithm (Turner et al., 2013). All chains were assessed to have converged with the Gelman-Rubin statistic, R-hat < 1.1. An individual participant's parameters were estimated as the mean of the corresponding participant-level posterior distributions.

To assess the fits of extracted model parameters in predicting behavior, we used the fitted computational parameters for each participant to obtain model-predicted choice rates and median reaction times (RTs). We assessed the model's ability to capture between-participant variability by correlating the observed and predicted choice rates and median RTs and then conducting a one-sample t-test on the choice and RT correlation coefficients.

Moreover, to assess the effects of experimental conditions on the estimated values for the hyper-mean parameters, we compared the overlap in posterior parameter distributions (called posterior probabilities) and converted the probabilities to two-tailed tests, with $P$-values used analogously to classical $p$-values. For example, to assess potential differences in non-decision time (*ndt*) estimates for the need and merit conditions of the social perception task, we evaluated the overlap between their respective posterior distributions. We inferred a significant difference between the conditions when the $P$-value from the two-tailed test was < 0.025 (i.e., $P < 0.05/2$), accounting for the two tails of the distribution (see Supplemental Note S5).

**Note S4.** Preprocessing of neuroimaging data.

*Preprocessing of neuroimaging data for the social perception and altruism task.*

Results included in this manuscript come from preprocessing performed using *fMRIPrep* 20.2.3 (Esteban, Markiewicz, et al. (2018); Esteban, Blair, et al. (2018); RRID:SCR_016216), which is based on *Nipype* 1.6.1 (Gorgolewski et al. (2011); Gorgolewski et al. (2018); RRID:SCR_002502).

*Anatomical data preprocessing.* A total of 1 T1-weighted (T1w) images were found within the input BIDS dataset. The T1-weighted (T1w) image was corrected for intensity non-uniformity (INU) with N4BiasFieldCorrection (Tustison et al. 2010), distributed with ANTs 2.3.3 (Avants et al. 2008, RRID:SCR_004757), and used as T1w-reference throughout the workflow. The T1w-reference was then skull-stripped with a *Nipype* implementation of the antsBrainExtraction.sh workflow (from ANTs), using OASIS30ANTs as target template. Brain tissue segmentation of cerebrospinal fluid (CSF), white-matter (WM) and gray-matter (GM) was performed on the brain-extracted T1w using fast (FSL 5.0.9, RRID:SCR_002823, Zhang, Brady, and Smith 2001). Brain surfaces were reconstructed using recon-all (FreeSurfer 6.0.1, RRID:SCR_001847, Dale, Fischl, and Sereno 1999), and the brain mask estimated previously was refined with a custom variation of the method to reconcile ANTs-derived and FreeSurfer-derived segmentations of the cortical gray-matter of Mindboggle (RRID:SCR_002438, Klein et al. 2017). Volume-based spatial normalization to two standard spaces (MNI152NLin6Asym, MNI152NLin2009cAsym) was performed through nonlinear registration with antsRegistration (ANTs 2.3.3), using brain-extracted versions of both T1w reference and the T1w template. The following templates were selected for spatial normalization: *FSL's MNI ICBM 152 non-linear 6th Generation Asymmetric Average Brain Stereotaxic Registration Model* [Evans et al. (2012), RRID:SCR_002823; TemplateFlow ID: MNI152NLin6Asym], *ICBM 152 Nonlinear Asymmetrical template version 2009c* [Fonov et al. (2009), RRID:SCR_008796; TemplateFlow ID: MNI152NLin2009cAsym],

*Functional data preprocessing.* For each of the BOLD runs (1 for the social perception task, 5 for the altruism task) found per participant (across all tasks and sessions), the following preprocessing was performed. First, a reference volume and its skull-stripped version were generated by aligning and averaging 1 single-band references (SBRefs). A B0-nonuniformity map (or *fieldmap*) was estimated based on two (or more) echo-planar imaging (EPI) references with opposing phase-encoding directions, with 3dQwarp Cox and Hyde (1997) (AFNI 20160207). Based on the estimated susceptibility distortion, a corrected EPI (echo-planar imaging) reference was calculated for a more accurate co-registration with the anatomical reference. The BOLD reference was then co-registered to the T1w reference using bbregister (FreeSurfer) which implements boundary-based registration (Greve and Fischl 2009). Co-registration was configured with six degrees of freedom. Head-motion parameters with respect to the BOLD reference (transformation matrices, and six corresponding rotation and translation parameters) are estimated before any spatiotemporal filtering using mcflirt (FSL 5.0.9, Jenkinson et al. 2002). BOLD runs were slice-time corrected using 3dTshift from AFNI 20160207 (Cox and Hyde 1997, RRID:SCR_005927). First, a reference volume and its skull-stripped version were generated using a custom methodology of *fMRIPrep*. The BOLD time-series (including slice-timing correction when applied) were resampled onto their original, native space by applying a single, composite transform to correct for head-motion and susceptibility distortions. These resampled BOLD time-series will be referred to as *preprocessed BOLD in original space*, or just *preprocessed BOLD*. The BOLD time-series were resampled into several standard spaces, correspondingly generating the following *spatially-normalized, preprocessed BOLD runs*: MNI152NLin6Asym, MNI152NLin2009cAsym. First, a reference volume and its skull-stripped version were generated using a custom methodology of *fMRIPrep*. Automatic removal of motion artifacts using independent component analysis (ICA-AROMA, Pruim et al. 2015) was performed on the *preprocessed BOLD on MNI space* time-series after removal of non-steady state volumes and spatial smoothing with an isotropic, Gaussian kernel of 6mm FWHM (full-width half-maximum). Corresponding “non-aggresively” denoised runs were produced after such smoothing. Additionally, the “aggressive” noise-regressors were collected and placed in the corresponding confounds file. Several confounding time-series were calculated based on the *preprocessed BOLD*: framewise displacement (FD), DVARS and three region-wise global signals. FD was computed using two formulations following Power (absolute sum of relative motions, Power et al. (2014)) and Jenkinson (relative root mean square displacement between affines, Jenkinson et al. (2002)). FD and DVARS are calculated for each functional run, both using their implementations in *Nipype* (following the definitions by Power et al. 2014). The three global signals are extracted within the CSF, the WM, and the whole-brain masks. Additionally, a set of physiological regressors were extracted to allow for component-based noise correction (*CompCor*, Behzadi et al. 2007). Principal components are estimated after high-pass filtering the *preprocessed BOLD* time-series (using a discrete cosine filter with 128s cut-off) for the two *CompCor* variants: temporal (tCompCor) and anatomical (aCompCor). tCompCor components are then calculated from the top 2% variable voxels within the brain mask. For aCompCor, three probabilistic masks (CSF, WM and combined CSF+WM) are generated in anatomical space. The implementation differs from that of Behzadi et al. in that instead of eroding the masks by 2 pixels on BOLD space, the aCompCor masks are subtracted a mask of pixels that likely contain a volume fraction of GM. This mask is obtained by dilating a GM mask extracted from the FreeSurfer’s *aseg* segmentation, and it ensures components are not extracted from voxels containing a minimal fraction of GM. Finally, these masks are resampled into BOLD space and binarized by thresholding at 0.99 (as in the original implementation). Components are also calculated separately within the WM and CSF masks. For each CompCor decomposition, the *k* components with the largest singular values are retained, such that the retained components’ time series are sufficient to explain 50 percent of variance across the nuisance mask (CSF, WM, combined, or temporal). The remaining components are dropped from consideration. The head-motion estimates calculated in the correction step were also placed within the corresponding confounds file. The confound time series derived from head motion estimates and global signals were expanded with the inclusion of temporal derivatives and quadratic terms for each (Satterthwaite et al. 2013). Frames that exceeded a threshold of 0.5 mm FD or 1.5 standardised DVARS were annotated as motion outliers. All resamplings can be performed with *a single interpolation step* by composing all the pertinent transformations (i.e. head-motion transform matrices, susceptibility distortion correction when available, and co-registrations to anatomical and output spaces). Gridded (volumetric) resamplings were performed using antsApplyTransforms (ANTs), configured with Lanczos interpolation to minimize the smoothing effects of other kernels (Lanczos 1964). Non-gridded (surface) resamplings were performed using mri_vol2surf (FreeSurfer).

Many internal operations of *fMRIPrep* use *Nilearn* 0.6.2 (Abraham et al. 2014, RRID:SCR_001362), mostly within the functional processing workflow. For more details of the pipeline, see the section corresponding to workflows in *fMRIPrep*'s documentation.

**Note S5.** Sanity check confirming enhanced perceptual sensitivity scores in task-relevant blocks of the social perception task.

In the social perception task, we estimated three sensitivity parameters ($S_{merit,}$ $S_{need}$, and $S_{control}$) independently for all three conditions (i.e., 9 sensitivity parameters total; 3 estimates x 3 conditions), irrespective of whether the judgment was task-relevant or not in the block. This allowed for the possibility that evidence relevant to a specific perceptual "quantity" might be sampled in contexts where it was not explicitly required and might influence judgments even when it was irrelevant. That said, we would expect larger perceptual sensitivity estimates in task-relevant settings (e.g., enhanced merit sensitivity in merit blocks of the social perception task). To confirm this assumption (which in essence serves as a manipulation check), we evaluated the effects of experimental conditions on the estimated sensitivity values for the hyper-mean parameters in our computational model of social perception. Specifically, we compared parameter posteriors and converted the probabilities to two-tailed tests.

Perceptual sensitivity estimates differed between the three conditions, as would be expected. In each of the blocks, the task-relevant sensitivity estimates (e.g., $S_{need}$ in the need condition) were significantly greater than they were in either of the other conditions (e.g., $S_{need}$ in the merit and factual control blocks; all Ps < 0.0001). Furthermore, $S_{control}$ did not differ between the need and merit conditions (P = 0.21), and $S_{merit}$ did not differ between the need and control conditions (P = 0.46). However, $S_{need}$ was significantly more positive in the merit block relative to the control conditions (P = 0.02).

Finally, participants exhibited a perceptual bias (overall drift *bias*) towards the "yes" response in the merit condition (suggesting that, on average, people tend to perceive others as deserving) but not in the need or control conditions (see Supplemental Table S2). The bias $w_0$ was significantly more negative in the need condition than in the merit condition (P = 0.002). The bias $w_0$ in the control condition was not significantly different from the need (P = 0.06) or merit conditions (P = 0.10) using 2-tailed statistical tests.

**Note S6.** Multivariate decoding of the inference condition in the social perception task.

We also performed multivariate decoding analysis on the brain data collected in the social perception task. We first estimated another General Linear Model (GLM) for each participant (using SPM12). The GLM was identical to the one used for the univariate analysis with two exceptions: first, the model was estimated on non-smoothed brain data. Second, instead of collapsing both task blocks per condition into one regressor of interest, the GLM modeled each block separately, yielding six regressors of interest (condition: merit, need, control x block: 1, 2).

The multivariate pattern analysis aimed to identify localized activation patterns that reliably decoded the perception condition (merit, need, control) in the social perception task. To ensure an unbiased analysis of the neural activation patterns throughout the whole brain, a "searchlight" approach was used (Haynes et al., 2007; Kriegeskorte et al., 2006). Given that this approach does not depend on a priori assumptions about informative brain regions or prior voxel selection, the problem of circular analysis (or "double dipping") can be avoided (Kriegeskorte et al., 2009). We implemented three separate decoding analyses: merit vs. control, need vs. control, merit vs. need. All three analyses used a similar analysis approach, which we illustrate using the example of the decoding of need vs. merit inferences below.

We used the same searchlight approach as in prior work (Tusche et al., 2016). For each participant, a sphere with a radius of 4 voxels was created around every voxel $v_i$ of the measured brain volume (Kahnt et al., 2011; Tusche et al., 2014; Wisniewski et al., 2015). For each sphere, we investigated whether the local pattern of activation predicted the inference condition (e.g., need vs. merit). For every task block, parameter estimates from the GLM (see above) were extracted for each of the N voxels in the sphere around voxel $v_i$ and transformed into an N-dimensional pattern vector. In total, for binary decoding analysis, we created four pattern vectors for each sphere (e.g., for a searchlight analysis decoding of need vs. merit, we extracted two block-wise pattern vectors for need, and two for merit). The first two of the pattern vectors (one per condition) were used for the training ("training dataset") of a linear support vector machine classifier with a fixed regularization parameter C = 1 (implemented using libSVM operated in MATLAB, https://www.csie.ntu.edu.tw/~cjlin/libsvm/). This provided the basis for the subsequent classification of the remaining two pattern vectors (one per condition) that were not used for the training. The procedure was then repeated by training on the last two pattern vectors and testing on the first two pattern vectors (yielding a two-fold cross-validation). The amount of condition-related information of the spatial activation pattern of each spherical cluster was represented by the average decoding accuracy across both cross-validation steps and was assigned to the central voxel $v_i$ of the cluster.

The described classification was performed for all spherical clusters created around every measured voxel, resulting in a three-dimensional map of average classification accuracies for each participant. These participant-specific accuracy maps were then spatially smoothed with an isotropic Gaussian kernel of 6 mm FWHM (full-width half-maximum). Finally, a standard second-level statistical analysis was performed to identify brain regions that allowed classifying the perceptual condition (e.g., whether individuals performed need or merit judgments) across participants ($n = 44$). More specifically, individual accuracy maps were submitted to a one-sample t-test and contrasted against chance level (as implemented in SPM12). Since the classification was based on two alternatives (e.g., merit vs. need), chance level was 50%. Only regions passing a stringent statistical threshold ($p < 0.001$ at the voxel level, $p < 0.05$ FWE corrected at the cluster level) and showing significant decoding accuracies above chance were considered relevant for information encoding (Haynes et al., 2007; Soon et al., 2008). Results for the multivariate decoding analyses are summarized in Supplemental Table S10.

Specifically, these additional analyses corroborated the involvement of the mentalizing network in the processing of merit (vs. control) and need (vs. control) inferences. Moreover, these supplemental analyses failed to identify multivariate activation patterns that reliably decoded need versus merit inferences in the social perception task ($p < 0.001$ at the voxel level, FWE corrected at the cluster level at $p < 0.05$). In other words, multivoxel activation patterns did not allow decoding whether participants were currently judging others' need or merit at this statistical level, suggesting common neural codes for both types of social perceptual judgments.

**Note S7. Brain responses in the altruism task and subsequent analyses.**

***General linear model (GLM) of neural responses during social choices.***

GLM2 examined how social contexts (recipient's need and merit) alter neural responses during altruistic choices. For each participant, we estimated a general linear model (GLM) using a canonical hemodynamic response function and a 128s high-pass cut-off filter to eliminate low-frequency drifts in the data. GLM2 estimated six regressors of interest for each of the five functional runs of the altruism task. Regressors of interest corresponded to the choice periods in each of the experimental cells of our 3 (partners' merit [high, low, unknown]) x 2 (partners' need [high, low]) factorial task design. Choice periods were defined by the onset of the choice screen and the response (button press) in that trial (Figure 1B). We also included several regressors of no interest: parametric modulators of choice periods by trial-wise decision values (1-4, "strong no" to "strong yes"), condition-wise regressors for missed trials (if applicable), six motion regressors and one regressor for the framewise displacement estimated during the preprocessing of the functional brain data, and session constants. For each participant, we created six condition-wise contrast images of choice periods against the implicit baseline.

We then estimated a full factorial design at the group level as implemented in SPM12 (http://www.fil.ion.ucl.ac.uk/spm). This allowed us to explore the neural correlates of the main effects of partner's need and merit that we observed on the behavioral level for the altruism task. We used the following statistical threshold for this group-level analysis: $p < 0.001$ at the voxel level, FWE corrected at the cluster level at $p < 0.05$ (matching the statistical threshold used for brain data from the social perception task).

In addition, we also set up several group level models to explore if the neural correlates of the main effects for need or merit in the altruism task covary with an individual's change in generosity (model free) or attribute weights (model-based) in response to partners' merit or need. We used participant-specific contrasts of interest (e.g., high merit – low merit) to set up a group level model (simple t-test as implemented in SPM12) and participant-specific change scores (e.g., generosity for a [high – low merit partners]) as covariates. We then tested if merit- (or need-) evoked changes in neural activations reflected the degree to which partner merit (or need) altered individuals' altruistic behavior. All whole brain analyses at the group level were thresholded at $p < 0.001$ at the voxel level, FWE corrected at the cluster level at $p < 0.05$.

***Neural underpinnings of merit- and need-evoked changes in altruistic choice***.

First, we aimed to identify brain regions that are differentially activated for [high vs. low merit] or [high vs. low need] conditions of the altruism task. There were no significant results for either merit or need ($p < 0.001$ at the voxel level, FWE corrected at the cluster level at $p < 0.05$). Second, we tested for brain activation that reflected individual differences in the responsiveness of our need or merit manipulation. Here, we used participant-specific change scores in generosity (model-free) and choice attributes (computational model-estimated attribute weights for \$self-, \$other, and fairness) as covariates of need- (or merit-) induced changes in brain activation at the group level. There were no significant results at our statistical threshold of $p < 0.001$ at the voxel level, FWE corrected at the cluster level at $p < 0.05$. Given the comparatively small sample size (n=28) due to COVID-related lockdowns, these null findings should be interpreted with caution.
